# CDKL5 regulates p62-mediated selective autophagy and host antiviral defense

**DOI:** 10.1101/2022.09.20.508746

**Authors:** Josephine W. Thinwa, Zhongju Zou, Emily Parks, Salwa Sebti, Kelvin Hui, Yongjie Wei, Vibha Singh, Greg Urquhart, Jenna L. Jewell, Julie K. Pfeiffer, Beth Levine, Tiffany A. Reese, Michael U. Shiloh

## Abstract

Virophagy, the selective autophagosomal engulfment and degradation of viral components, is crucial for antiviral immunity. However, the mechanisms leading to viral antigen recognition and autophagy induction remain poorly understood. Here, we identify a novel kinase, Cyclin-dependent kinase-like 5 (CDKL5), as an essential regulator of virophagy. Deletion of CDKL5 or abrogation of its kinase activity reduced virophagy of Sindbis virus (SINV), a neurotropic RNA virus, and increased intracellular accumulation of SINV capsid proteins and cellular cytotoxicity. Mechanistically, through direct phosphorylation of the selective autophagy receptor p62, CDKL5 promoted formation of p62 inclusion bodies that bound capsid. Loss of CDKL5 disrupted the capsid-p62 interaction, and a p62 phosphomimetic mutant rescued the interaction. *CDKL5* knockout mice demonstrated increased neuronal cell death after SINV infection and enhanced lethality after infection with several human viruses. Overall, these findings identify a cell-autonomous innate immune mechanism for autophagy activation to clear toxic viral capsid aggregates during infection.

## Introduction

Upon infection, host cells become virus production factories. Accumulation of large quantities of viral proteins overwhelms cellular degradative capabilities, leading to protein aggregation and cell death (Ahmad et al., 2018). Host cell survival thus depends not only on preventing viral protein synthesis, but also on eliminating viral debris. Autophagy is a cell-autonomous process for cellular sequestration of unwanted cytoplasmic components such as damaged organelles, protein aggregates and invading pathogens into double membrane vesicles for subsequent lysosomal degradation (Levine and Kroemer, 2019; Mizushima and Levine, 2020). Cargo uptake occurs either non-selectively or selectively based on cargo recognition by specific autophagy receptors. The selective autophagic degradation of infectious cargo is called xenophagy. Viral xenophagy, or simply virophagy, targets viral antigens and particles to the autophagosome for degradation and plays an important role in cell-autonomous innate immunity. Several selective autophagy receptors participate in xenophagy: p62/sequestosome 1 (SQSTM1), OPTN, NDP52, NBR1, and TAX1BP1 (Viret et al., 2021). However, to date, the specific host factors involved in regulating receptor-cargo recognition to initiate virophagy remain poorly understood.

Neurons, a non-renewable cell population, rely heavily on virophagy for protection against viral infection. For example, virophagy is critical for the antiviral response and host survival during SINV infection (Liang et al., 1998; Orvedahl et al., 2010). SINV is a positive sense RNA virus that causes murine viral encephalitis in a manner analogous to other arthropod-borne viruses. Overexpression of an essential autophagy gene, *Beclin 1*, decreases SINV replication and apoptosis of neuronal cells and enhances mouse survival after lethal intracerebral infection (Liang *et al*., 1998). Conversely, genetic disruption of another essential autophagy gene, *Atg5*, causes SINV capsid protein accumulation in both cultured cell lines and murine brain tissue with associated neuronal cell death, despite no impact on viral replication (Orvedahl *et al*., 2010). Thus, virophagy plays a cytoprotective role in neurons in part by preventing virus-induced cell death (Ahmad *et al*., 2018).

Given the involvement of autophagy in numerous cellular functions, identifying host factors that regulate the specific recognition of virus antigens for virophagy could be exploited for the creation of host-directed antiviral interventions. Recently, a genome-wide image-based siRNA screen in HeLa cells infected with SINV or herpes simplex virus type 1 (HSV-1) identified over 200 genes, including the serine/threonine kinase Cyclin-dependent kinase-like 5 (CDKL5), as candidate early regulators of virus-induced autophagy (Dong et al., 2021). We chose to further investigate the potential autophagy function of CDKL5 as it is widely expressed, with highest expression in neurons. Our results identify CDKL5 as a key initiator of autophagy through its phosphorylation of the canonical autophagy receptor p62 to facilitate subsequent engulfment of viral capsid aggregates. As CDKL5 functions in neuronal axon outgrowth, dendritic morphogenesis, and synaptogenesis, and human *CDKL5* mutations result in a severe neurodevelopmental disorder through a poorly understood mechanism (Bahi-Buisson and Bienvenu, 2012; Rusconi et al., 2008; Zhu and Xiong, 2019), these findings have implications for understanding both the molecular mechanisms of virophagy and the mechanisms by which CDKL5 regulates neuronal development.

## Results

### Basal and Virus-Induced Autophagy Requires CDKL5 in a Kinase-Dependent Manner

To investigate whether CDKL5, as identified on the genome-wide siRNA screen (Dong *et al*., 2021), impacts autophagy during virus infection, we generated two CDKL5-deficient clones (CDKL5 KO) in HeLa cells stably expressing GFP-LC3 using CRISPR/Cas9 (Figure 1A and 1B). As expected, SINV infection of parental HeLa (WT)/GFP-LC3 led to a significant increase in GFP-LC3 puncta (Figure 1A and 1C). In contrast, compared to WT HeLa/GFP-LC3 cells, both CDKL5 KO HeLa/GFP-LC3 clones had fewer GFP-LC3 puncta under basal conditions and after autophagy induction via SINV infection (Figure 1A and 1C). Conversion of LC3-I to the autophagosome-associated LC3-II by lipidation also was reduced in CDKL5 KO HeLa cells compared with WT cells after SINV infection (Figures 1D, 1E and S1). Reconstitution of CDKL5 KO cells with WT CDKL5 rescued the autophagy defect (Figure 1A-1E). Deletion of *CDKL5* did not impact autophagy induction by other canonical stimuli including amino-acid starvation and mTOR inhibition based on GFP-LC3 puncta formation and levels of endogenous LC3-II (Figure S2A-S2C, related to Figure 1). Taken together, these data suggest that CDKL5 regulates both basal and SINV-induced autophagy, but not other canonical forms of autophagy induced by nutrient or energy stress.

**Figure 1.**
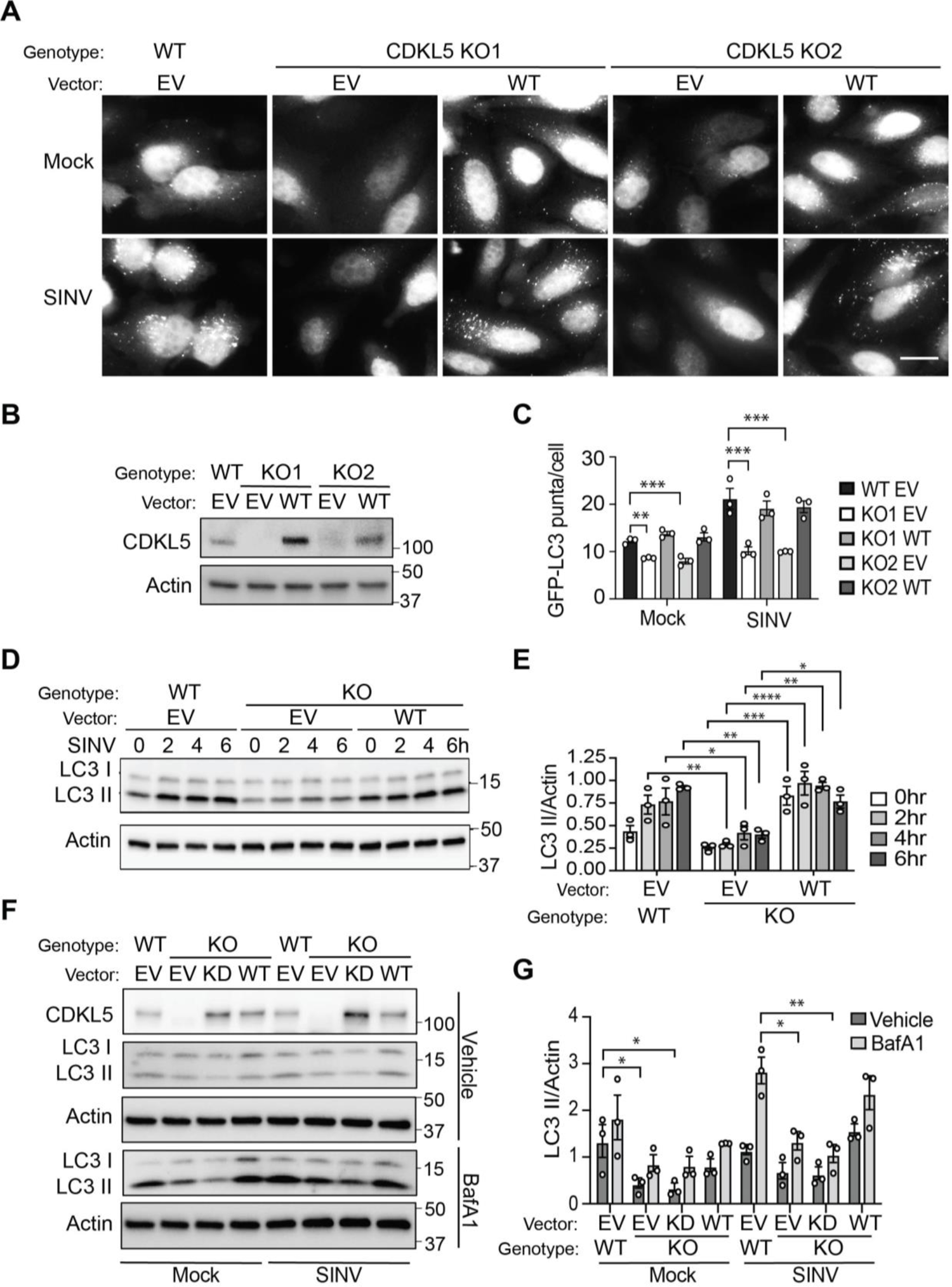
CDKL5 is necessary for basal and virus induced autophagy WT CDKL5/GFP-LC3 and two CDKL5 KO/GFP-LC3 HeLa clones (KO1 and KO2) were reconstituted with CDKL5 or empty vector (EV). (A) Representative fluorescent micrographs of GFP-LC3 puncta (autophagosomes) in mock cells or cells infected with WT SINV at a multiplicity of infection (MOI) of 10 for 6 h. Scale bar represent 20 μm. (B) Western blot of CDKL5 expression in WT and KO cells reconstituted with WT CDKL5 or empty vector (EV). (C) Quantification of GFP-LC3 puncta with bars representing mean ± SEM of triplicate samples with at least 100 cells per sample. Statistical analysis by one-way ANOVA with Dunnett’s test for multiple comparisons (\*\**p* < 0.01 and \*\*\**p* < 0.001). Results are representative of three independent experiments. (D) Western blot of LC3 conversion over a 6h time course comparing WT and CDKL5 KO HeLa cells reconstituted with WT CDKL5 or EV infected with SINV (MOI=10). E) Quantification of LC3-II signal normalized to actin displayed as mean ± SEM from three independent experiments; significance was assessed using one-way ANOVA with Sidak’s multiple comparison test (\**p* < 0.05, \*\**p* < 0.01, \*\*\**p* < 0.001, and \*\*\*\**p* < 0.0001). (F, G) Western blot of LC3 conversion (F) and quantification by densitometry (G) comparing WT and CDKL5 KO cells reconstituted with EV, WT or a kinase dead (KD) CDKL5, infected with SINV (MOI=10; 6 h) and treated with either DMSO (vehicle) or BafA1 for 2 h prior to harvesting cells. Statistical analysis performed by one-way ANOVA with Dunnett’s test for multiple comparisons (\**p* < 0.05 and \*\**p* < 0.01).

We next tested if the kinase activity of CDKL5 is required for autophagy induction. To that end, we complemented the CDKL5 KO HeLa cells with a kinase dead mutant through a point mutation (K42R) that targets the ATP-binding site within the catalytic domain known to be clinically relevant in humans and associated with CDKL5 deficiency disorder (Bahi-Buisson et al., 2008; Bertani et al., 2006). Unlike complementation of CDKL5 KO HeLa cells with WT CDKL5, the kinase dead CDKL5^K42R^ mutant (KD CDKL5) failed to rescue the levels of lipidated LC3-II at baseline or after SINV infection (Figure 1F and 1G). Notably, treating cells with the lysosomal inhibitor bafilomycin A1 (BafA1) caused an increase in LC3-II levels at baseline and after SINV infection in all genotypic backgrounds indicating that autophagic flux was maintained in CDKL5 deficient cells (Figure 1F and 1G).

### CDKL5 Controls Viral Protein Clearance and Cell Survival

Because control of SINV infection requires virophagy (Orvedahl *et al*., 2010), we investigated the impact of CDKL5 deficiency on viral capsid accumulation. We infected WT and CDKL5 KO HeLa cells with recombinant SINV expressing an mCherry-capsid fusion protein for direct visualization of capsid via fluorescence microscopy (Figure 2A) and flow cytometry (Figure 2B and 2C). Loss of CDKL5 resulted in robust intracellular accumulation of mCherry-capsid proteins (Figures 2A-2C). Reconstitution of CDKL5 KO cells with WT CDKL5 significantly decreased capsid levels, whereas reconstitution with KD CDKL5 did not rescue the capsid accumulation phenotype observed in CDKL5-deficient cells (Figure 2D and 2E). Thus, we concluded that CDKL5 through its kinase activity regulates capsid accumulation after SINV infection.

**Figure 2.**
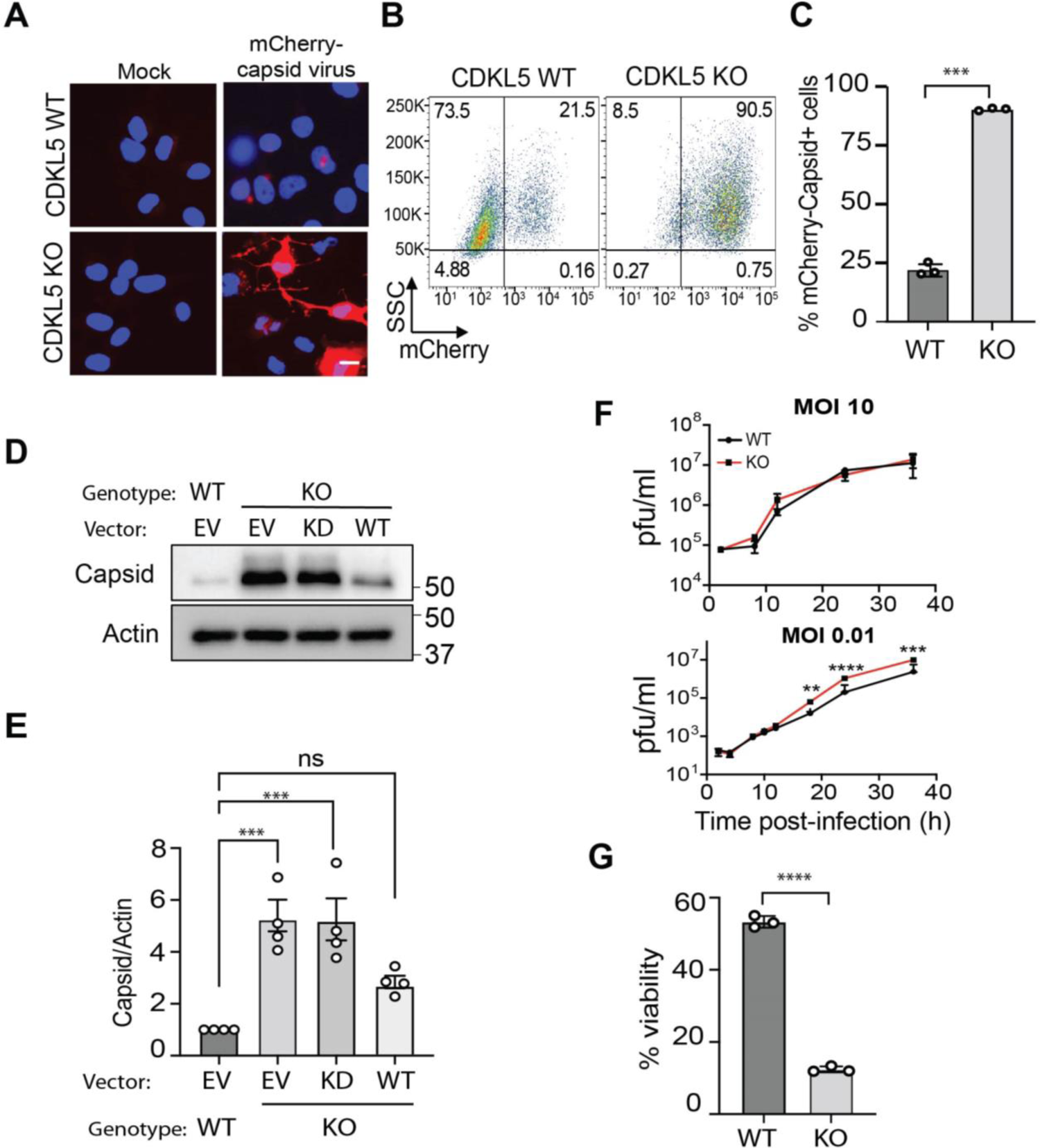
CDKL5-deficient cells accumulate SINV capsid protein and have increased cell death (A-C) WT and CDKL5 KO HeLa cells were mock infected or infected with SINV/mCherry-capsid (MOI=10) for 24 hours. (A) Representative fluorescent micrographs of mCherry-capsid; Scale bar represent 20 μm. (B) Representative flow cytometry plots of mCherry-capsid positive cells and (C) quantification of three independent replicates. Bars represent mean ± SEM. *p* values were determined by unpaired t-test (\*\*\**p* < 0.001). (D,E) WT and CDKL5 KO HeLa cells reconstituted with EV, KD or WT *CDKL5* were infected with SINV/mCherry-capsid (MOI=10, 24h) and (D) mCherry-capsid accumulation detected by Western blot using anti-SINV capsid antibody. (E) Quantification of capsid/actin ratio from four independent experiments. p values analyzed by one-way ANOVA with Dunnett’s test for multiple comparisons (\*\*\**p* < 0.001). (F) High MOI growth curve analysis (MOI 10) and low MOI multi-step growth curve analysis (MOI 0.01) of WT and CDKL5 KO HeLa cells infected with SINV/mCherry-capsid virus. Progeny viruses were quantified by plaque assays. Bars represent mean ± SD of three experimental replicates and *p* values were determined by two-way ANOVA with Sidak’s multiple comparison test (\*\**p* < 0.01, \*\*\**p* < 0.001, and \*\*\*\**p* < 0.0001). (G) WT and CDKL5 KO HeLa cells were infected with SINV/mCherry-capsid virus (MOI=10, 24h) and cell viability determined by CellTiter Glo assay. Bars on graph represent mean ± SEM from three independent experiments; *p* values were determined by unpaired two-tailed t-test (\*\*\*\**p* < 0.0001).

One possible explanation for the increased capsid accumulation is increased viral replication in CDKL5 KO cells, though prior work demonstrated that impaired removal of viral proteins via virophagy did not directly correlate with significantly increased viral replication (Dong *et al*., 2021; Orvedahl *et al*., 2010; Orvedahl et al., 2011; Sumpter et al., 2016). In agreement with previous reports, we observed no difference in SINV replication between WT and CDLK5 KO cells when we performed a high multiplicity of infection (MOI, 10) growth curve analysis, and only a modest increase in replication in CDKL5 KO cells in a low MOI (0.01) multi-cycle growth curve analysis (Figure 2F).

Viral protein clearance via autophagy plays an additional cytoprotective role during virus infection by mitigating the toxicity of viral protein aggregates that accumulate in the cell (Ahmad *et al*., 2018). To determine the impact of CDKL5 on cell survival during viral infection, we infected WT and CDKL5 KO HeLa cells with SINV/mCherry-capsid. Loss of CDKL5 markedly decreased cell viability after SINV infection compared to WT cells, suggesting that CDKL5 promotes cell survival during SINV infection (Figure 2G).

### Primary Cortical Neurons Deficient in CDKL5 Have Impaired Autophagy After Virus Infection

Because neurons robustly express CDKL5 and serve as a physiologically relevant cell-type for SINV infection (Hector et al., 2016), we sought to determine the autophagy function of CDKL5 in mouse primary cortical neurons. We isolated cortical neurons from CDKL5 WT/GFP-LC3 and CDKL5 KO/GFP-LC3 mouse embryos, infected the neurons with SINV and measured autophagy induction by quantifying GFP-LC3 puncta using fluorescence microscopy. While neurons of both genotypes had increased GFP-LC3 puncta in response to amino acid starvation, only WT neurons demonstrated increased autophagy induction after SINV infection (Figure 3A and 3B). In addition, CDKL5-deficient neurons increasingly had higher levels of capsid post infection (Figure 3C and 3D) suggesting that CDKL5 plays a role in the mitigation of capsid accumulation in neurons.

**Figure 3.**
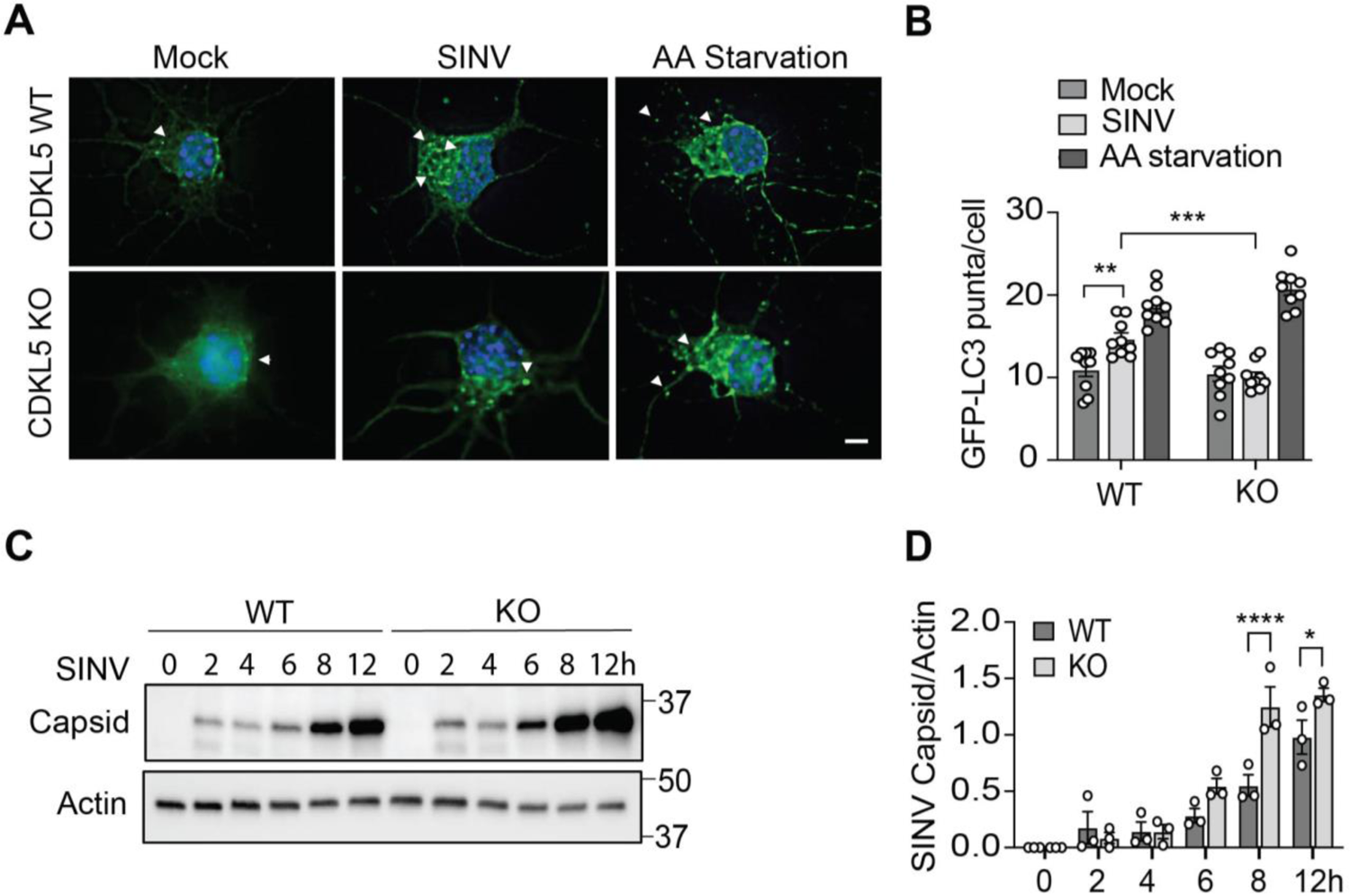
Virophagy but not amino acid starvation-induced autophagy is dependent on CDKL5 in mouse primary cortical neurons Primary cortical neurons were isolated from littermate CDKL5 WT/GFP-LC3 and CDKL5 KO/GFP-LC3 embryos and either maintained in normal media, WT SINV infected (MOI = 10, 8h) or treated with EBSS (amino acid (AA) starvation) medium for 2h. (A) Representative fluorescent micrographs. Arrows point to representative GFP-LC3 puncta. Scale bar represents 10 μm. (B) Quantification of GFP-LC3 puncta. Bars are mean ± SEM from triplicate samples (at least 50 neurons per sample counted) from four independent embryos per genotype. *p* values were determined by one-way ANOVA with Dunnett’s test for multiple comparison (\*\**p* < 0.01 and \*\*\**p* < 0.001). (C-D) Western blot of capsid accumulation detected using anti-SINV capsid antibody in *CDKL5*^WT^/GFP-LC3 and *CDKL5*^KO^/GFP-LC3 neurons infected with WT SINV (MOI =10, time points as indicated) with (D) quantification of SINV capsid/actin ratio from three independent experiments. Bars represent mean ± SEM; scale bar 10 μm. Statistical analysis performed by one-way ANOVA test with Sidak’s multiple comparison test (\**p* < 0.05 and \*\*\**p* < 0.0001).

### CDKL5 Regulates Viral Capsid Clearance Independent of Viral Replication

Since CDKL5 may restrict SINV capsid accumulation through several mechanisms that impact the synthesis and clearance of capsid, we tested the impact of CDKL5-deficiency on viral capsid clearance independent of viral replication. To that end, we used a UV-inactivated SINV capable of cellular uptake but not replication. To detect autophagy induction with UV-inactivated SINV, we treated cells with the equivalent of an MOI of 500. We pulsed WT and CDKL5 KO HeLa cells with UV-inactivated SINV and after removing extracellular UV-inactivated virus, quantified cellular capsid clearance over time. Though both WT and KO cells internalized UV-inactivated SINV equally, capsid was cleared by 2-3h in WT cells but persisted in CDKL5 KO cells (Figure 4A and 4B). To confirm that capsid clearance is autophagy dependent, we determined the rate of UV-inactivated SINV capsid clearance by autophagy deficient HeLa ATG7 KO cells (Figure S3A, related to Figure 4). Absence of ATG7 resulted in delayed capsid clearance compared to WT cells (Figure S3A and 3B, related to Figure 4) indicating that autophagy plays a role in capsid clearance. Notably, SINV capsid could still be cleared by HeLa ATG7 KO cells, albeit more slowly, suggesting that other mechanisms can reduce SINV capsid accumulation in the absence of autophagy. We further investigated the role of autophagy in capsid clearance by treating WT and CDKL5 KO cells with either BafA1 to prevent lysosomal acidification and thus autophagic flux (Klionsky et al., 2021) or PIK-III, a Class III phosphatidylinositol-3 kinase (PI3KC3, also known as VPS34) inhibitor, which inhibits PI3KC3 C1 and C2 complexes essential for autophagosome membrane nucleation and lysosome fusion (Dowdle et al., 2014). Whereas treatment of CDKL5 KO cells with both inhibitors had no significant effect on capsid levels, treatment of WT cells showed significantly more capsid after 1 h of the chase period compared to vehicle (Figure 4C-4E). Similarly, WT cells pulsed with UV-inactivated virus had a greater accumulation of lipidated LC3 in the presence of BafA1, whereas PIK-III treated cells did not accumulate lipidated LC3 (Figure 4C and 4F). Taken together, these findings demonstrate that CDKL5 mediates the efficient clearance of capsid from UV-inactivated SINV in an autophagy-dependent manner.

**Figure 4.**
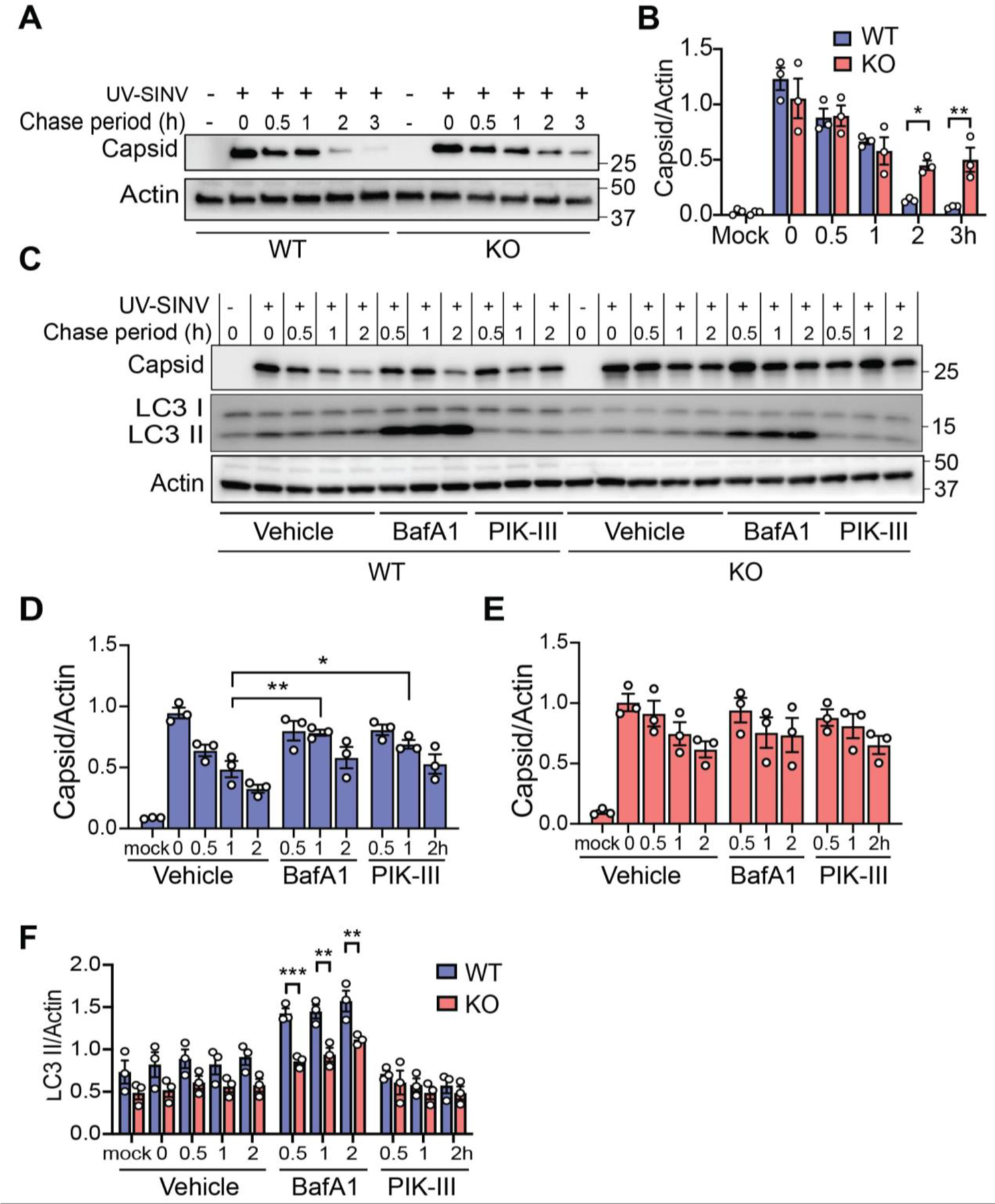
Loss of CDKL5 delays autophagy-mediated clearance of capsid in cells pulsed with non-replicating SINV (A-B) WT and CDKL5 KO HeLa cells exposed to UV-inactivated WT SINV for 1 h, washed, and lysates harvested at the indicated time points. (A) Western blot of capsid accumulation and (B) quantification of capsid/actin ratio by densitometry. Bars are mean ± SEM from three independent experiments. *p* values were determined by one-way ANOVA test with Sidak’s multiple comparison (\**p* < 0.05 and \*\**p* < 0.01). (C-F) WT and CDKL5 KO HeLa cells exposed to UV-inactivated SINV for 1 h, treated with DMSO (vehicle), BafA1 (100 μM) or PIK-III (5 μ*M*) and cell lysates analyzed by (C) Western blot for capsid and CDKL5 accumulation and LC3 conversion. (D) Quantification of capsid/actin ratio in WT cells. (E) Quantification of capsid/actin ratio in CDKL5 KO cells. (F) Quantification of LC3 II normalized to Actin. (D-F) Bars are mean ± SEM from three independent experiments. *p* values in (D-E) were determined one-way ANOVA with Dunnett’s test for multiple comparisons (\**p* < 0.05 and \*\**p* < 0.01), and in (F) by one-way ANOVA test with Sidak’s multiple comparison test (\**p* < 0.05, \*\**p* < 0.01, and \*\*\**p* < 0.001).

### CDKL5 Promotes Association of Viral Capsid with p62

Because CDKL5 deficient cells demonstrated defective autophagy induction and delayed SINV capsid clearance, we hypothesized that CDKL5 impacts the interaction of SINV capsid with the selective autophagy receptor p62 to initiate virophagy. p62 interacts with SINV capsid in HeLa cells to facilitate capsid clearance (Orvedahl *et al*., 2010). To investigate the role of CDKL5 in the capsid-p62 interaction, we infected WT and CDKL5 KO HeLa cells with recombinant SINV expressing HA-tagged capsid and performed co-immunoprecipitation (coIP) of p62 after HA pull-down. As expected, endogenous p62 in WT cells co-immunoprecipitated with HA-capsid (Figure 5A). Surprisingly, CDKL5 also co-immunoprecipitated with HA-capsid (Figure 5A). However, in the absence of CDKL5, we did not detect p62 after HA-capsid immunoprecipitation despite higher levels of capsid in the cell lysates (Figure 5A) and similar p62 transcript and baseline protein levels (Figure S4, related to Figure 5). In addition, baseline levels of the core selective autophagy receptors NBR1, p62, NDP52, OPTN, and TAX1BP1 were similar when comparing WT and KO cells (Figure S4, related to Figure 5). To further characterize the association between p62 with SINV capsid, we infected WT and CDKL5 KO HeLa cells (Figure 5B and 5C) and primary cortical neurons isolated from CDKL5 WT and KO mice (Figure S5, related to Figure 5) with SINV/mCherry-capsid and performed immunofluorescence colocalization studies at an early time-point before cells lacking CDKL5 became saturated with capsid. We observed a greater number of colocalized mCherry-capsid/p62 puncta in WT compared to CDKL5 KO cells (Figures 5B, 5C and S5). Taken together, these data suggest that CDKL5 facilitates the interaction of p62 with SINV capsid.

**Figure 5.**
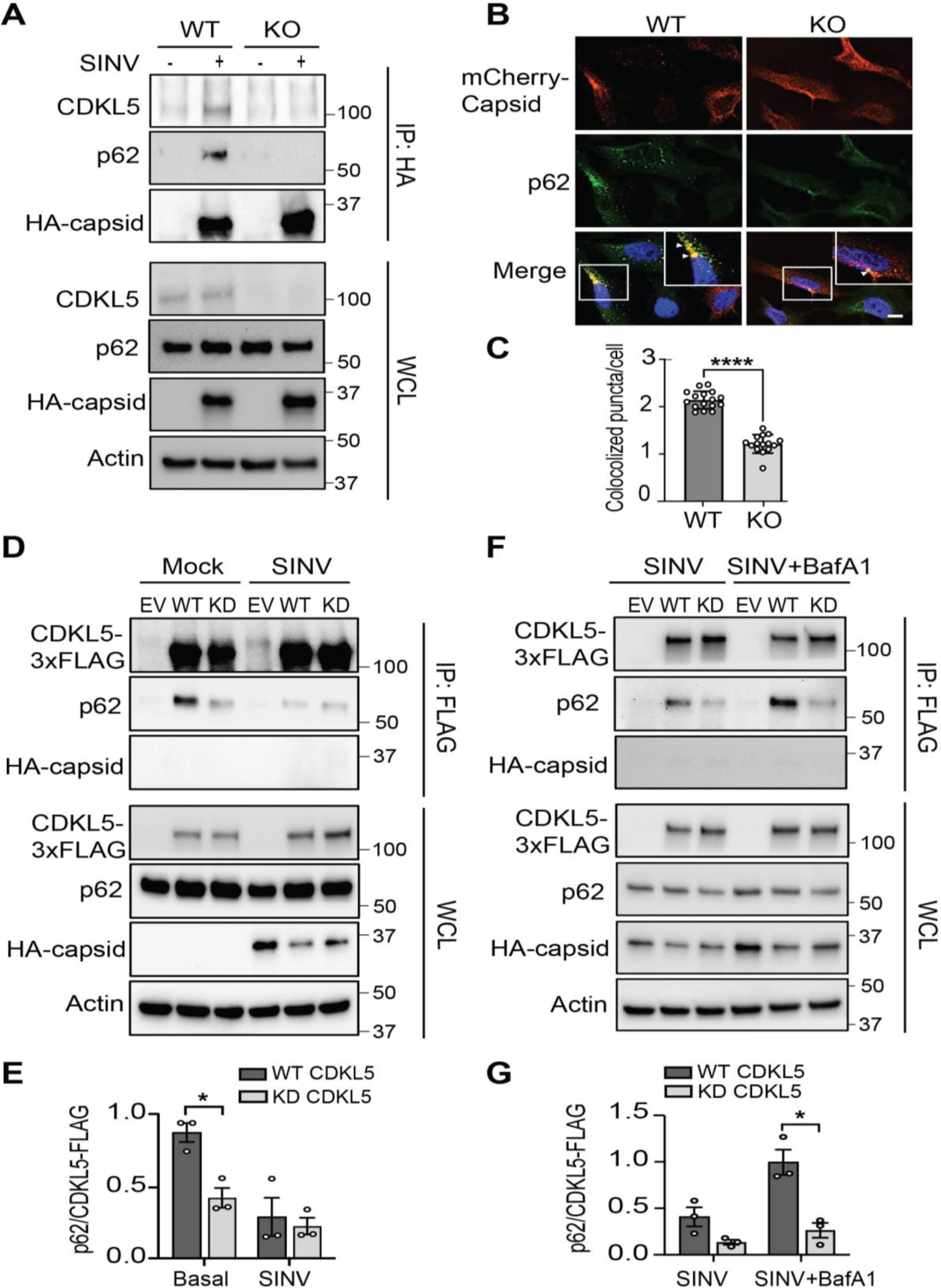
CDKL5 is necessary for the interaction of SINV capsid with p62. (A) Immunoblot of endogenous CDKL5, p62 and HA-capsid after HA-capsid coimmunoprecipitation from *CDKL5*^WT^ and *CDKL5*^KO^ HeLa cells infected with SINV/HA-capsid (MOI = 10, 7h). Results are representative of three independent experiments. (B, C) WT and CDKL5 KO HeLa cells were infected with SINV/mCherry-capsid (MOI=10, 7h) then stained with antibody to p62. (B) Representative immunofluorescence of p62 and mCherry-capsid with (C) quantification of colocalization. Bars are mean ± SEM of mCherry-capsid^+^/p62^+^ puncta with at least 50 cells in triplicate samples from 4 independent experiments analyzed. Arrowheads denote representative colocalized capsid/p62 puncta. Scale bar, 10 μm. \*\*\*\**p* < 0.0001 determined by unpaired two-tailed *t*-test. (D, E) CDKL5 KO HeLa cells reconstituted with EV, WT or KD CDKL5 fused to a C-terminal 3xFLAG were then infected with SINV/HA-capsid (MOI = 10, 7h). (D) Immunoblot with anti-HA, anti-p62 and anti-CDKL5 antibodies after CDKL5-3xFLAG immunoprecipitation with anti-FLAG antibody. (E) Quantification of p62 that coimmunoprecipitated with WT or KD CDKL5-FLAG normalized to FLAG. Bars are mean ± SE from three independent experiments. *p* value was determined by one-way ANOVA with Sidak’s multiple comparison test (\**p* < 0.05). (F, G) CDKL5 KO HeLa cells reconstituted as in (D) were infected with SINV infected cells treated with either vehicle (DMSO) or BafA1 (100 μM) for 2h. (F) Immunoblot performed as in (D) and (G) quantification of p62 coimmunoprecipitation as in (E). Bars are mean ± SE from three independent experiments. *p* value was determined by one-way ANOVA with Sidak’s multiple comparison test (\**p* < 0.05).

Because the kinase activity of CDKL5 was necessary for virophagy (Figure 1), we next tested if CDKL5 kinase activity impacts the interaction of CDKL5 with capsid. We expressed WT CDKL5-FLAG and KD CDKL5-FLAG in CDKL5 KO HeLa cells and performed anti-FLAG immunoprecipitation in mock-infected or SINV/HA-capsid infected cells. Neither WT nor KD CDKL5-FLAG immunoprecipitation resulted in detectable HA-capsid co-immunoprecipitation (Figure 5D, 5F). However, in uninfected cells we detected p62 after immunoprecipitation of WT CDKL5-FLAG and to a lesser extent after immunoprecipitation with the KD CDKL5 (Figure 5D and 5E). Furthermore, when cells were infected with SINV, the coimmunoprecipitation of p62 by WT CDKL5-FLAG was greatly diminished (Figure 5D and 5E). Blocking autophagic flux with BafA1 increased the amount of p62 that co-immunoprecipitated with WT CDKL5-FLAG but did not increase p62 co-immunoprecipitation by KD CDKL5-FLAG. These data suggest that during SINV infection, CDKL5/p62 interaction induces the autophagic clearance of p62 (Figure 5F and 5G). Taken together, our data demonstrate that CDKL5 interacts with p62, and the kinase activity of CDKL5 is necessary for autophagic degradation of p62 during viral infection.

### CDKL5 phosphorylates p62 to promote binding to capsid

Because the kinase activity of CDKL5 was necessary for virophagy induction (Figure 1) and interaction of CDKL5 with p62 during SINV infection of HeLa cells (Figure 5), we hypothesized that CDKL5 directly phosphorylates p62. To test this, we performed in vitro phosphorylation assays using recombinant CDKL5 and p62 proteins. As expected (Muñoz et al., 2018), in the absence of p62, CDKL5 underwent autophosphorylation, thus serving as a positive control for its catalytic activity (Figure 6A). While p62 alone did not autophosphorylate, combining CDKL5 with p62 resulted in phosphorylation of p62 (Figure 6A). An in vitro phosphorylation time course showed that CDKL5 autophosphorylation, which leads to autoactivation of the kinase, preceded the phosphorylation of p62 thus providing further evidence that p62 is a substrate of CDKL5 (Figure S6, related to Figure 6).

**Figure 6.**
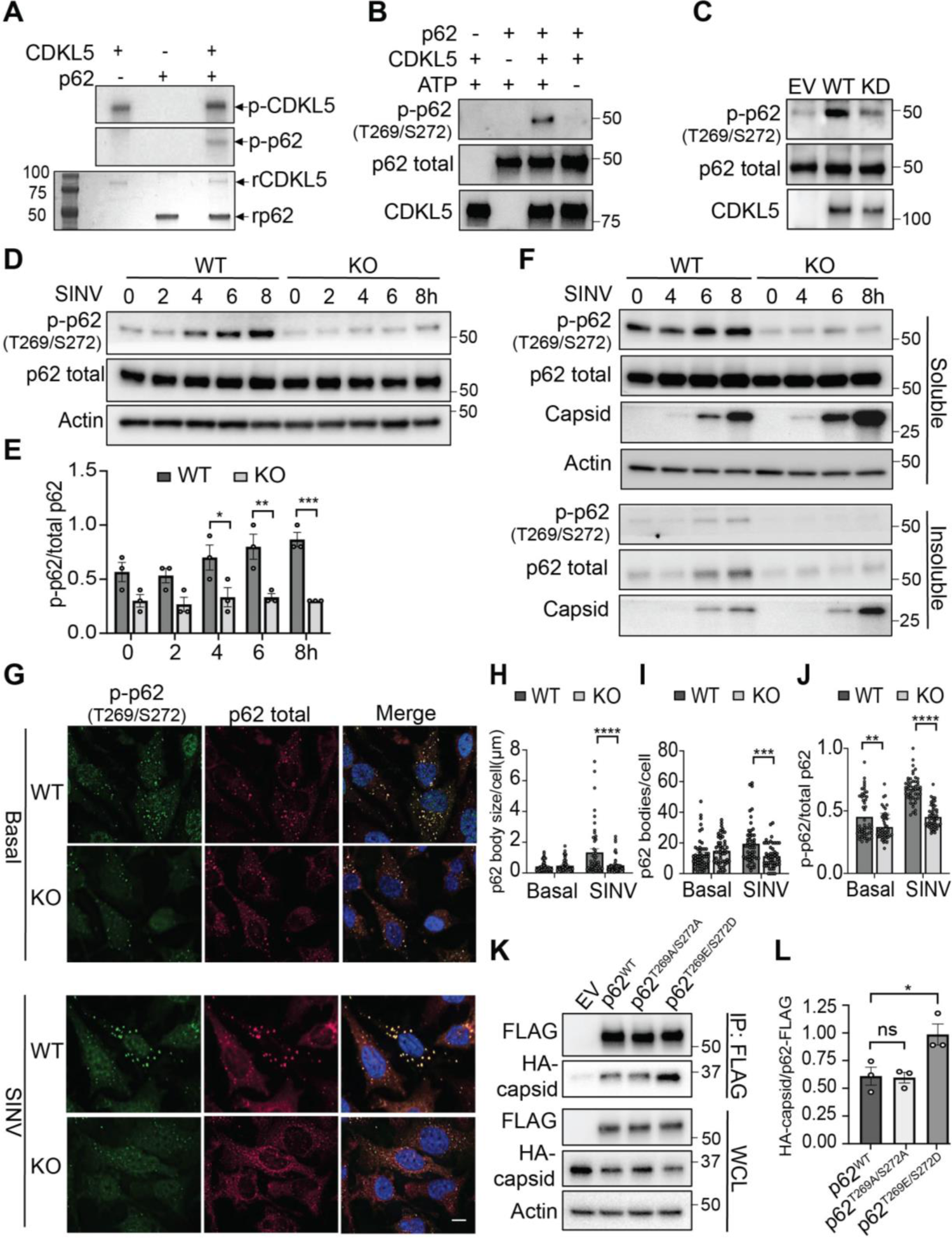
CDKL5 phosphorylates p62 at Thr269/Ser272 to impact binding to capsid. (A) In vitro kinase assay with recombinant CDKL5 (1-498 aa) and p62 using [^32^P] ATP. *Top* Phosphorylation of p62 and autophosphorylation of CDKL5 detected after a 30-minute reaction on ^32^P-autoradiography. The *bottom* panel shows Coomassie Brilliant Blue staining of the gel. (B) In vitro kinase assay with recombinant CDKL5 (1-498 aa) and p62 using non-radioactive ATP and detection of p62 phosphorylation by western blot using anti-T269/S272 phosphospecific antibody. Blot representative of three independent experiments. (C) EV, WT and KD CDKL5-FLAG were purified from HeLa cell lysates by immunoprecipitation and used in an in vitro kinase assay as in (B) with recombinant p62 as substrate. Phosphorylation of T269/S272 residues on p62 was detected by Western blot as in (B). Blot representative of three independent experiments. (D) WT and CDKL5 KO HeLa cells infected with SINV (MOI=10) were lysed directly in Laemmli buffer and total lysates analyzed for levels of phospho-T269/S272 p62 by western blot using anti-p62 and anti-T269/S272 phosphospecific antibodies. Blot representative of three independent experiments. (E) Quantification of phosphoT269/S272 normalized to total p62. Bars are mean ± SEM. *p* value was determined by one-way ANOVA with Sidak’s multiple comparison test (\**p* < 0.05, \*\**p* < 0.01, and \*\*\**p* < 0.001). (F) WT and CDKL5 KO HeLa cells were infected with SINV. At various time points after infection, lysates were prepared and fractionated into soluble and insoluble fractions in RIPA buffer containing 1% NP-40. The segregation of proteins into soluble versus insoluble fractions was detected by Western blotting using anti-capsid, anti-p62 and anti-phospho p62 (T269/S272) antibodies. Blot is representative of three independent experiments. (G-J) WT and CDKL5 KO HeLa cells were infected with SINV (MOI=10, 8h) or left uninfected, stained with anti-p62 or anti-phospho T269/S272 p62 antibodies for fluorescence microscopy. (G) Representative fluorescent micrographs are shown with analysis of the (H) size, (I) count and (J) ratio of phospho-T269/S272 p62 to total p62 bodies per cell quantified using Fiji software. 50 cells per condition were analyzed. Bars are mean ± SEM. Scale bar, 10 μm. *p* values were determined by one-way ANOVA with Sidak’s multiple comparison test (\*\**p* < 0.01, \*\*\**p* < 0.001, and \*\*\*\**p* < 0.0001). (K-L) CDKL5 KO HeLa cells stably expressing either EV, WT p62, p62 with alanine substitution (T269A/S272A) or phosphomimetic p62 (T269E/S272D) tagged with 3X-FLAG were infected with SINV/HA-capsid virus (MOI 10; 8h), cell lysates prepared and then subjected to immunoprecipitation using FLAG antibody. Capsid was detected by probing with anti-HA antibody, and p62 variants were detected with anti-FLAG antibody. Representative blot of three independent experiments. (L) Quantification of capsid coimmunoprecipitating with p62-FLAG analyzed by densitometry and normalized to FLAG signal. Bars are mean ± SEM from three independent experiments. *p* value was determined by one-way ANOVA with Dunnett’s multiple comparison test (\**p* < 0.05).

We next sought to identify a site on p62 that CDKL5 phosphorylates. p62 has multiple established phosphorylation sites (Gubas and Dikic, 2022; Kumar et al., 2022), and antibodies are commercially available for the most common sites. We initially hypothesized that CDKL5 phosphorylates p62 at S403 because phosphorylation of this site increases the binding affinity of p62 to ubiquitinated cargo during selective autophagy including bacterial xenophagy (Matsumoto et al., 2015; Matsumoto et al., 2011; Pilli et al., 2012). To test our hypothesis, we used an in vitro phosphorylation assay to achieve CDKL5 phosphorylation of p62 but found no detectable S403 phosphorylation (Figure S6B, related to Figure 6). In contrast, Tank Binding Kinase-1(TBK-1), a kinase known to phosphorylate p62 at S403 (Matsumoto et al., 2011; Pilli et al., 2012), led to detectable phospho-p62 (Figure S6B, related to Figure 6).

Since our data demonstrated that the absence of CDKL5 reduced colocalization of p62 with capsid within large punctate structures (Figure 5B and 5C), we hypothesized that CDKL5 phosphorylation influences the ability of p62 to cluster around capsid. Notably, during proteotoxic stress caused by proteosome inhibition, phosphorylation of p62 at T269/S272 by p38 MAPK facilitates formation of p62 aggresomes that sequester misfolded proteins to initiate autophagosome biogenesis (Zhang et al., 2018). To test whether CDKL5 phosphorylates p62 at T269/S272, we performed in vitro kinase assay and probed with a phospho-antibody specific for phosphorylated T269/S272. We detected phospho-p62 (T269/S272) only in the presence of both CDKL5 and ATP (Figure 6B). As an additional line of evidence, we affinity-purified WT CDKL5-FLAG or KD CDKL5-FLAG expressed in CDKL5 KO HeLa cells and tested the ability of WT and KD CDKL5 to phosphorylate recombinant p62 in vitro. We found greater phosphorylation of p62 in the presence of WT CDKL5 when compared to EV-FLAG and KD CDKL5-FLAG (Figure 6C). To test the impact of SINV infection on CDKL5-dependent T269/S272 phosphorylation of endogenous p62, we infected WT and CDKL5 KO HeLa cells with SINV and assessed p62 T269/S272 phosphorylation over time. Wild type HeLa cells demonstrated not only higher basal phospho-T269/S272 p62 compared to CDKL5 KO cells (Figure 6D and 6E, compare time=0), but also a significantly greater increase in T269/S272 p62 phosphorylation after infection with SINV (Figure 6D and 6E). Taken together, these findings demonstrate that CDKL5 phosphorylates p62 at T269/S272 and SINV infection induces CDKL5-dependent T269/S272 phosphorylation.

We next sought to determine whether p62 aggregation was defective in the absence of CDKL5 during SINV infection. Whereas we previously assessed p62 levels in total cell lysates (Figure 6D), to detect p62 aggresome formation we fractionated cell lysates to RIPA-soluble and -insoluble fractions from SINV-infected WT and CDKL5 KO HeLa cells, as nascent p62 aggresomes are expected to partition into the insoluble fraction (Sarraf et al., 2020). While soluble total p62 levels were equal between SINV-infected WT and CDKL5 KO HeLa cells, p62 accumulated to a greater extent in the insoluble fraction isolated from WT than KO cells (Figure 6F). Furthermore, phospho-p62 T269/S272 was detectable in the insoluble fraction of WT but not KO cells (Figure 6F). Interestingly, CDKL5 KO cells overall had more capsid in both the soluble and insoluble fractions suggesting that while aggregation of p62 depends on CDKL5, capsid aggregation is independent of CDKL5 (Figure 6F). We further examined the clustering of p62 into large intracellular bodies by immunofluorescence imaging and found that compared to SINV-infected CDKL5 KO cells, WT cells had significantly larger and greater numbers of p62 bodies with proportionately more phospho-T269/S272 contained within these bodies (Figure 6G-6J). These findings indicate that CDKL5 deficient cells are defective in the formation and incorporation of phospho-p62 T269/S272 into p62 aggresomes.

Since the aggregation of p62 functions in cargo recruitment to autophagosomes (Wurzer et al., 2015; Zaffagnini et al., 2018), we predicted that if we overexpressed a phosphomimetic mutant p62^T269E/S272D^ (Zhang *et al*., 2018) in CDKL5 KO cells, we could rescue p62 interaction with capsid. To that end, we stably expressed FLAG-tagged versions of WT p62, p62 that cannot be phosphorylated (p62^T269A/S272A^) and the phosphomimetic mutant (p62^T269E/S272D^) in CDKL5 KO HeLa cells. We assessed the interaction between p62 and HA-capsid through coimmunoprecipitation using FLAG antibody. WT p62 and p62^T269A/S272A^ bound similar levels of capsid (Figure 6K and 6L). In contrast, the phosphomimetic mutant p62^T269E/S272D^-FLAG bound significantly more capsid (Figure 6K and 6L), suggesting that CDKL5 phosphorylation of these residues facilitates the interaction between p62 and capsid.

### CDKL5 is required for host defense against diverse viruses

We next investigated the impact of CDKL5 on viral antigen clearance and neuronal cell death *in vivo*. We intracranially infected neonatal WT or *CDKL5*^KO^ mice with SINV and collected their brains post infection to assess the role of CDKL5 in controlling viral pathology and replication. Using immunohistochemistry to detect viral capsid and TUNEL staining to detect apoptotic cells, we observed that mice lacking CDKL5 demonstrated increasing levels of capsid and apoptotic cells in the brain over a 7-day time course, while WT mice showed an opposite trajectory with both clearance of capsid and apoptotic cells by day 7 after infection (Figure 7A-7D). Despite these divergent histopathological findings, both mouse genotypes had similar SINV replication based on viral titers on days 1 and 4 indicating that increased capsid levels on day 4 in *CDKL5*^KO^ mice is not due to increased viral replication (Figure 7C and 7E). By day 7, *CDKL5^KO^* mice demonstrated higher viral titers compared to WT mice (Figure 7E). In conclusion, these data support the hypothesis that CDKL5 initiates autophagic clearance of viral antigens and maintenance of tissue viability after viral infection.

**Figure 7.**
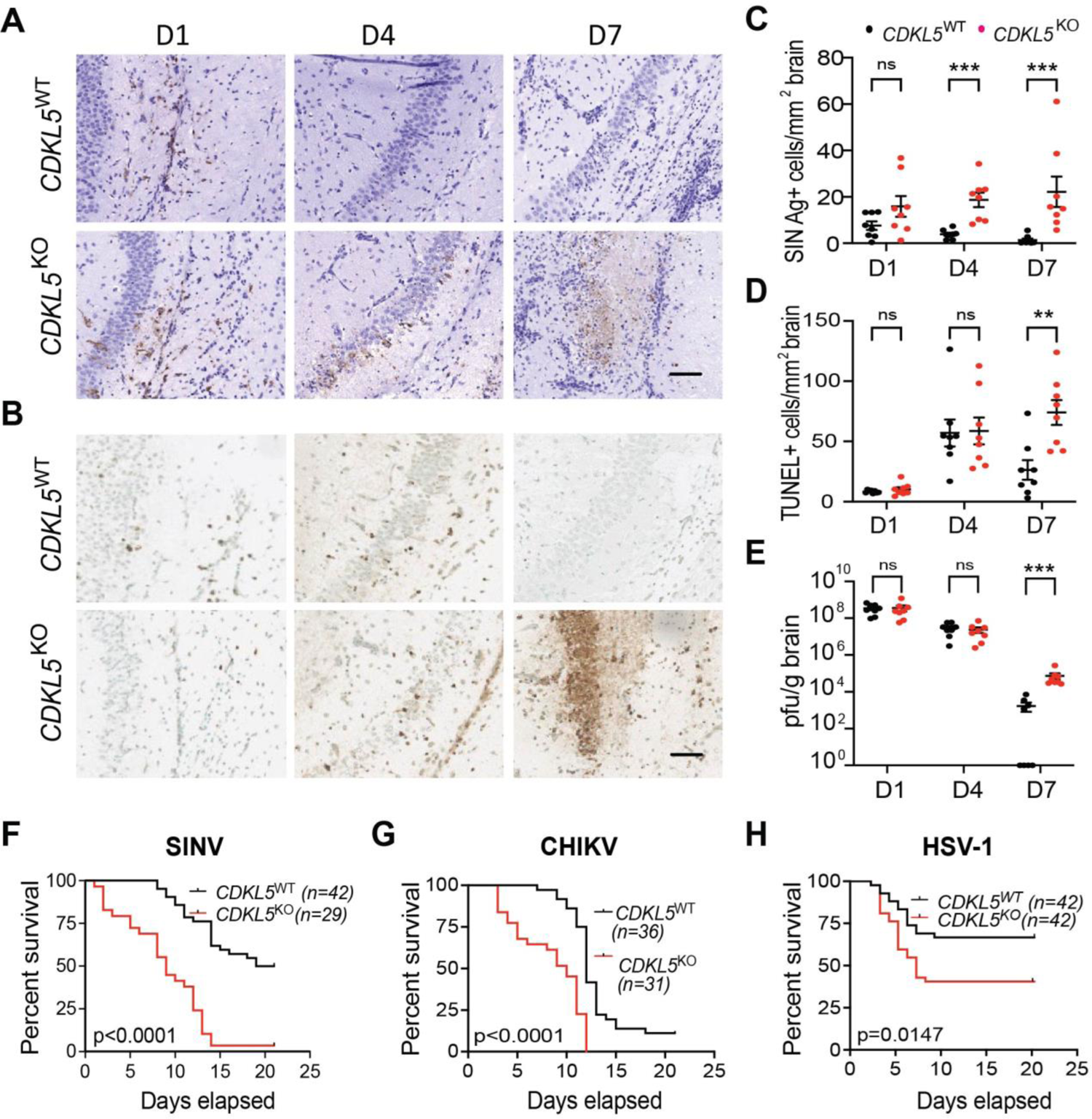
CDKL5 is required for host defense against virus infection in mice (A-E) Seven-day-old *CDKL5*^WT^ and *CDKL5*^KO^ mice were infected intracerebrally (i.c.) with SINV strain dsTE12Q (1 x 10^3^ plaque forming units (pfu)). Brains from 8 mice per genotype were harvested at the indicated days post infection. (A) The presence of SINV capsid positive cells was determined by immunohistochemistry. Representative light micrographs of capsid by immunohistochemistry. Scale bars, 100 μm. (B) Identification of apoptotic cells in brains sections using TUNEL assay. Representative light micrographs. Scale bars, 200 μm. (C) Quantification of capsid-positive cells per section of brain. Bars represent mean ± SEM of 7-8 mice per group. *P* value determined by Mann-Whitney U test (\*\*\**p* < 0.001). (D) Quantification of TUNEL-positive area per brain section. Bars represent mean ± SEM of 7-8 mice per group. \*\**p* < 0.01 was determined by Mann-Whitney U test. (E) Viral titers from brain homogenates processed at the indicated days post infection. Bars represent mean ± SEM of 7-8 mice per group. Statistical analysis by Mann-Whitney U test (\*\*\**p* < 0.001). (F-H) Survival of 7-day-old *CDKL5*^WT^ and *CDKL5*^KO^ mice infected (F) i.c. with SINV or (G) subcutaneously with CHIKV (1 x 10^5^ pfu) and (H) 8-10-week-old *CDKL5*^WT^ and *CDKL5*^KO^ mice infected i.c. with HSV-1ΔBBD (5 x 10^4^ pfu). Number of mice per genotype and *p* value (log rank) are indicated on the graph.

To determine if CDKL5 has a more generalized role in host antiviral immune responses, we used three murine viral infection models previously shown to require autophagy. We infected mice with either SINV or with two additional neurotropic viruses with significant impact on human health: herpes simplex type-1 strain HSV-1ΔBBD, a DNA virus lacking the autophagy inhibitory activity of ICP34.5 (Orvedahl et al., 2007) or chikungunya virus (CHIKV) (Joubert et al., 2012), a single-stranded RNA virus in the same genus as SINV. Intracranial infection with SINV or HSV-1ΔBBD causes fatal murine encephalitis in neonatal or adult mice, respectively, while subcutaneous infection with CHIKV causes lethality of neonatal mice (Dong *et al*., 2021; Orvedahl *et al*., 2007; Orvedahl *et al*., 2010; Orvedahl *et al*., 2011; Sumpter *et al*., 2016). Compared to WT mice, *CDKL5^KO^* mice were markedly more susceptible to SINV infection and succumbed to infection more rapidly than WT mice (Figure 7F). Similarly, *CDKL5*^KO^ mice were more susceptible to infection with CHIKV and HSV-1ΔBBD compared to wild-type littermates (Figure 7G-7H), suggesting that CDKL5 has a generalized function in host antiviral defense.

## Discussion

Here, we report a heretofore unknown function for CDKL5 as a regulator of selective autophagy of viruses. Our results demonstrate that CDKL5 has a cytoprotective role in SINV infected cultured cells by facilitating the autophagic clearance of viral proteins. Mechanistically, we identify the selective autophagy receptor p62 as a novel substrate of CDKL5 and provide evidence that CDKL5 facilitates the interaction between SINV capsid with p62. In addition to the in vitro findings, we show that CDKL5 is an essential factor in host antiviral defense against diverse viruses by demonstrating that *CDKL5*^KO^ mice have enhanced mortality after infection with SINV, CHIKV and HSV-1.

A role for CDKL5 in autophagy has not previously been described. CDKL5 is a serine/threonine kinase important in neurodevelopment, as loss of function mutations in humans result in a severe X-linked neurodevelopmental disorder called CDKL5 deficiency disorder (CDD) (Katayama et al., 2020). By using a combination of genetically ablated cell lines and primary cortical neurons, we demonstrate that CDKL5 is required for the induction of autophagy during SINV infection. Moreover, a human disease associated CDKL5 kinase dead mutant failed to rescue the autophagy defect, indicating that its autophagy regulatory function depends on its kinase activity. These findings raise the possibility that defective autophagy contributes to the neurological pathogenesis of CDD.

The molecular mechanisms underlying the pathogenesis of CDD and the phosphorylation targets of CDKL5 are not well defined. Notably several neurological pathologies ranging from neurodegenerative to neurodevelopmental disorders are associated with defective autophagy (Levine and Kroemer, 2019). Failure of selective autophagy to clear protein aggregates or damaged mitochondria has been recognized as a pathologic mechanism of neurodegeneration in Alzheimer’s disease, Parkinson’s disease, and Amyotrophic Lateral Sclerosis (Fleming et al., 2022). Likewise, dysfunctional mitochondrial clearance (mitophagy) has been implicated as a possible cause of the neurodevelopmental disorder Rett syndrome, which shares similar features as CDD (Crivellari et al., 2021; Kadam et al., 2019; Sbardella et al., 2017). Thus, selective autophagy, mediated in part by CDKL5, is critical for normal human development, neuronal homeostasis, and the protective response to cellular stressors like organellar damage and infection. In selective autophagy, the discriminate targeting of cargo to the growing autophagosome depends on autophagy receptors (Gubas and Dikic, 2022). Receptor-cargo recognition followed by recruitment of the cargo to autophagosomes involves numerous regulatory proteins particularly kinases and ubiquitin ligases making post-translational modifications (PTMs) on autophagy receptors and cargo. While the core autophagy receptor p62 is known to interact with SINV capsid (Orvedahl *et al*., 2010), the precise molecular trigger that initiates engagement of p62 with capsid was not known. Here, we demonstrate that CDKL5 functions as a regulator of p62 cargo recognition by enhancing the interaction between p62 and SINV capsid through phosphorylation of p62 at T269/S272.

p62 undergoes a variety of PTMs, with phosphorylation being the most common and abundant PTM, to direct its many cellular functions ranging from regulating signaling pathways to delivering cargo to autophagosomes and proteosomes (Gubas and Dikic, 2022). The wide array of p62 cellular functions is attributed to the ability of p62 to assemble into different oligomeric structures ranging from homo-dimers to large cytoplasmic inclusion bodies (Gubas and Dikic, 2022). Formation of large p62 inclusion bodies results in association with ubiquitinated cargo and its trafficking to forming autophagosomes (Bjørkøy et al., 2005; Wurzer *et al*., 2015; Zaffagnini *et al*., 2018). In the absence of CDKL5, phosphorylation of T269/S272 is markedly diminished and p62 less efficiently forms large cytoplasmic inclusion bodies or interacts with capsid. The expression of a phosphomimetic mutant of p62^T269E/S272D^ in CDKL5 KO cells rescues capsid binding suggesting that phosphorylation of these residues plays an important role in the clustering of p62 with capsid cargo. This result also highlights the critical importance of this phosphorylation event in the context of infection.

How does p62 T269/S272 phosphorylation by CDKL5 affect cell autonomous innate immunity against viruses? One possible explanation relates to the role of p62 in response to the accumulation of misfolded proteins. Proteotoxic stress due to proteosome inhibition induces p38 MAPK-mediated p62 phosphorylation at T269/S272 to activate the coalescing of p62 microaggregates into large perinuclear bodies to eliminate misfolded proteins through autophagy (Zhang *et al*., 2018). Viral replication also leads to rapid accumulation of large quantities of viral antigens mimicking proteotoxic stress. Therefore, the accumulation of viral protein aggregates may trigger CDKL5-mediated phosphorylation of p62 at T269/S272, activating autophagic clearance of accumulating viral antigens. Remaining critical questions are what specific signal activates CDKL5 to phosphorylate p62 and whether CDKL5 phosphorylates other residues on p62.

In our animal studies, *CDKL5*^KO^ mice were more susceptible to infection compared to their WT littermates after infection with several genetically diverse viruses. Notably, p62 serves as a virophagy receptor for capsid proteins from several viruses including: the double-stranded RNA virus avibirnavirus, the positive sense single-stranded RNA viruses coxsackievirus, CHIKV, poliovirus and foot-and-mouth disease virus, and the DNA virus human cytomegalovirus (Berryman et al., 2012; Judith et al., 2013; Li et al., 2020; Mohamud et al., 2019; Zimina et al., 2021; Zimmermann et al., 2021). The divergent life cycles of these viruses suggest that shared signals such as proteotoxic stress and aggregation of large quantities of viral antigens facilitate capsid recognition by p62. Thus, beyond SINV, CDKL5 may serve as a master regulator of p62-mediated autophagic clearance of viral antigens from multiple classes of viruses.

Histopathological analysis of SINV-infected murine brains revealed that in addition to capsid accumulation, loss of CDKL5 resulted in neuronal cell death. This result correlates with the observed enhanced cytotoxicity of infected CDKL5 KO HeLa cells. These findings align with other previously published studies showing that disrupting autophagy results in accumulation of viral protein aggregates and increases virus-induced neuronal cell death during SINV, West Nile Virus and HSV-1 infections (Kobayashi et al., 2012; Kobayashi et al., 2020; Orvedahl *et al*., 2010; Sumpter and Levine, 2016). The precise mechanism of how autophagy mitigates virus-induced cell death remains to be fully investigated.

In summary, our results reveal a previously unidentified role of CDKL5 as a regulator of virophagy and demonstrate its importance in host immunity to neurotropic DNA and RNA viruses. Both *in vivo* and *in vitro*, CDKL5 prevents the toxic accumulation of viral antigens and promotes cell survival during virus infection. The identification of CDKL5 and its kinase activity as critical for the interaction between SINV capsid and a key selective autophagy receptor p62 suggests novel targets for therapeutics aimed at increasing anti-viral autophagy.

## Methods

### Cell culture and reagents

HeLa cells were purchased from American Type Culture Collection (ATCC) and all clones were cultured in Opti-MEM I Reduced Serum Medium supplemented with 5% FBS, 100 units/mL penicillin and 100 μg/mL streptomycin, all purchased from Thermo Fisher Scientific. Cell authentication was performed at ATCC Cell Line Authentication Service. Two clones of HeLa *CDKL5* CRISPR knockout cell lines were generated through CRISPR/Cas9 based modification at the Genome Engineering and iPSC Center (GEiC) at Washington University School of Medicine (St. Louis, MO). Two guide RNAs were used: 5’-TACCTTCACCTACAACCCCANGG and 5’-TTTGAGATCCTTGGGGTTGTNGG to induce a double-strand break to exon 2. Knock-out clones underwent targeted deep-sequencing to assess for the presence of insertions and out-of-frame indels. To generate HeLa cells stably expressing GFP-LC3, cells were transfected with pIRES.GFP-LC3.puro plasmid followed by puromycin selection (1μg/ml). Lentiviruses expressing pLenti-C-Myc-DDK-IRES-Neo (OriGene, empty vector) or vector with encoded WT CDKL5 (NM_001323289.2), KD CDKL5, WT CDKL5-3xFLAG, KD CDKL5-3xFLAG, WT p62-3xFLAG, p62^T269A/S272A^-3xFLAG, or p62^T269E/S272D^-3x-FLAG were used to transduce CDKL5 KO HeLa cells followed by selection with geneticin (500ng/ml) (Thermo Fisher, 10131027). All genes were synthesized as geneblocks (IDT) and their sequences confirmed. ATG7 KO HeLa cells have been previously described (Selleck et al., 2015) and were a gift from Herbert Virgin. Vero cells used for plaque assays were cultured in Dulbecco’s Modified Eagle Medium (DMEM) (Thermo Fisher Scientific) supplemented with 10% FBS, 100 units/mL penicillin and 100 μg/mL streptomycin.

### Mouse strains

C57BL/6J and *CDKL5*^KO^ mice (Wang et al., 2012) backcrossed more than ten generations to C57BL/6J were obtained from The Jackson Laboratory. *CDKL5*^KO^ (JAX #021967) mice were additionally backcrossed for 3 generations with C57BL/6J breeders to achieve backcrossing to >12 generations. GFP-LC3 transgenic mice have been previously described (Mizushima et al., 2004), and were crossed with *CDKL5*^KO^ mice. All mouse strains were housed in a pathogen-free facility with 12-hour light/dark cycle and fed *ad libitum*. Age of mice used for specific experiments is indicated in the figure legends. Mouse experiments were reviewed and approved by the Institutional Animal Care and Use Committee at the University of Texas Southwestern and followed the eighth edition of the Guide for the Care and Use of Laboratory Animals. The University of Texas Southwestern is accredited by the American Association for Accreditation of Laboratory Animal Care (AAALAC).

### Preparation of cultured primary cortical neurons

To generate mouse cortical neurons, *CDKL5*^+/-^ female and *CDKL5*^+/y^ male or *CDKL5*^+/-^ female and *CDKL5*^+/y^/GFP-LC3^+/+^ male mice were mated and the brains from littermate offspring harvested at embryonic day 15 (E15). The cortex was dissected from the rest of the brain under a dissecting microscope. Cortical cells were dissociated in 2.5% Trypsin (Thermo Fisher Scientific) and DNase I (Sigma), and resuspended in DMEM supplemented with 10% FBS, 100 units/mL penicillin and 100 μg/mL streptomycin. Cells were then plated on Nunc Lab-Tek II 8-well glass chamber slides (Thermo Fisher Scientific) or 12-well dishes coated overnight with 0.1% polyethyleneimine (PEI) from Sigma. After 2 h, the medium was replaced with Neurobasal medium supplemented with B27 (Invitrogen), 2 mM glutamine (Thermo Fisher Scientific), and 5 μM Cytarabine (AraC) (Sigma) for 24h to suppress growth of non-neuronal cells, at which point fresh medium without AraC was added to culture neurons for 7 days.

### Antibodies

The following antibodies were used: rabbit anti-LC3B (Novus Biologicals, NB100-2220), guinea pig anti-p62/SQSTM1 (C-terminus) (Progen, GP62-C), mouse anti-SQSTM1 (Abnova, H00008878-M01), rabbit anti-phospho-SQSTM1/p62 (Thr269/Ser272) (Cell Signaling Technology, 13121), rabbit anti-phospho-SQSTM1/p62 (Ser403) (Cell Signaling Technology, 39786), mouse anti-actin antibody (C4) HRP (Santa Cruz Biotechnology (SCBT), sc-47778 HRP), mouse anti-CDKL5 (D-12) antibody (SCBT, sc-376314), rabbit anti-ATG7 polyclonal antibody (Sigma-Aldrich A2856), rabbit anti-TBK1/NAK antibody (Cell Signaling Technology, 3013), rabbit anti SINV capsid antibody (Gift from Diane Griffin) (Jackson et al., 1988), rabbit anti-HA tag (C29F4) antibody (Cell Signaling Technology, 3724), mouse anti-NBRI antibody, clone 6B11 (Abnova, H00004077-M01), rabbit anti-NDP52 antibody (Cell Signaling Technology, 60732S), rabbit anti-OPTN antibody (Proteintech, 10837-1-AP), rabbit anti-TAX1BP1 antibody (4H2L18) (Thermo Fisher, 702840), mouse anti-Flag M2 antibody (Sigma, F1804), donkey anti-rabbit IgG HRP-conjugate species-adsorbed (Millipore Sigma, AP182P), goat anti-mouse IgG (H+L) HRP (Millipore Sigma, AP308P), donkey anti-mouse IgG (H+L) highly cross-adsorbed secondary antibody Alexa Fluor 488 (Thermo Fisher Scientific, A21202), donkey anti-rabbit IgG (H+L) highly cross-adsorbed secondary antibody Alexa Fluor 488 (Thermo Fisher Scientific, A-21206), and donkey anti-rabbit IgG (H+L) highly cross-adsorbed secondary antibody Alexa Fluor 594 (Thermo Fisher Scientific, A21207).

### Viral strains

Sindbis virus strain SVIA (ATCC) was derived from a low-passage isolate of the wild-type SIN strain AR339 (Taylor et al., 1955). Recombinant SINV strain dsTE12Q engineered with a double subgenomic promoter was previously described (Liang *et al*., 1998). Recombinant SINV strain dsTE12Q.mCherry-capsid (SINV.mCherry-Capsid) was previously described (Orvedahl *et al*., 2010). For the generation of dsTE12Q.HA-capsid virus, a geneblock fragment “CATCTGACTAATACTACAACACCACCACCATGTACCCGTATGATGTTCCGGATTACGCTGGCTATCCCTACGACGTGCCCGACTATGCCGGGTACCCCTATGACGTCCCAGACTACGCA AATAGAGGATTCTTTAACATGCTCGGC” containing a partial capsid sequence and the in-frame 3xHA coding sequence was synthesized (IDT). The dsTE12Q recombinant vector was linearized by PCR with primers “AATAGAGGATTCTTTAACATGCTCGGC” and “GGTGGTGGTGTTGTAGTATTAGTCAGATG” immediately after the capsid start codon. The linearized vector and geneblock were then mixed to create the dsTE12Q.HA-Capsid recombinant SINV vector using NEBuilder^®^ HiFi DNA Assembly Master Mix per manufacturer’s instructions (New England BioLabs, E2621S). The mutant herpes simplex virus type 1 (HSV-1) strain (HSV-1ΔBBD) was previously described (Orvedahl *et al*., 2007). The chikungunya virus (CHIKV) strain 06-021 was a gift of Deborah J. Lenschow, Washington University. Viral stocks were propagated and titrated by plaque assays in either Vero or BHK-21 cells. The Sindbis virus strain SVIA was used in all infection studies with HeLa cells and primary cortical neurons unless otherwise specified in the figure legends.

### Infection with UV-inactivated SINV

To generate UV-inactivated virus, SINV strain SVIA equivalent to MOI of 500 was irradiated for 7 minutes using a Stratalinker UV Crosslinker 1800 (Stratagene). Absence of infectious particles was confirmed through plaque assay on Vero cells. After UV-inactivation, HeLa cells were exposed to virus particles for 1 h and then washed three times to remove extracellular or loosely bound virus before fresh culture media was added. Lysates were harvested for Western blot analysis at timed intervals to chase the clearance of capsid protein. To block autophagy induction and flux, 5 μM of PIK-III inhibitor (Selleck Chemicals, S7683) or 100nM of Bafilomycin A1 (Sigma-Aldrich B1793) were added 30 minutes after cells were exposed to UV-inactivated SINV.

### Reconstitution of HeLa *CDKL5^KO^* cells with CDKL5 and p62 alleles

Wild type CDKL5, WT CDKL5-3xFlag, KD CDKL5, KD CDKL5-3xFlag, WT p62-3xFlag, p62^T269A/S272A^−3xFlag, and p62 ^T269E/S272D^-3x Flag were cloned into pLenti-C-Myc-DDK-IRES-Neo vector (Origene) using NEBuilder^®^ HiFi DNA Assembly Master Mix per manufacturer’s instructions (New England BioLabs, E2621S). To generate the lentiviruses, the vectors were transfected into Phoenix cells (ATCC) together with packaging vectors pCMVR8.91 (Zufferey et al., 1997) and pMDG (Naldini et al., 1996). Cell culture supernatants were collected on day 2 and 3, filtered with a 0.45 μm filter (EMD Millipore) and then used to infect WT or CDKL5 KO HeLa cells in the presence of 8 μg/ml polybrene (SCBT, sc-134220). After 6 h, virus-containing media was exchanged with fresh culture media. After 48 h, 500 μg/ml of geneticin (500ng/ml) (Thermo Fisher, 10131027) was added for 7 days for selection and continued at 100 μg/ml for an additional 14 days prior to SINV experiments.

### Assays for autophagy assessment

To determine the induction of autophagy in HeLa cells and primary cortical neurons (Klionsky *et al*., 2021), the numbers of autophagosomes as represented by GFP-LC3 puncta per cells were counted by an observer blinded to the condition or cell genotype by fluorescence microscopy. In addition, LC3-I to LC3-II conversion was determined by western blot using anti-LC3B antibody. For virophagy experiments, cells were infected with SINV strains MOI of 10 in Opti-MEM I Reduced Serum Medium supplemented with 5% FBS (culture medium) for 1h then exchanged with fresh culture media or maintained in culture media only (mock). For amino-acid starvation, cells were treated for either 1h (GFP-LC3 microscopy assays) or 3 h (LC3 I/II western blots) with Earle’s Balanced Salt Solution (EBSS, Thermo Fisher Scientific) at 37℃ and for mTOR inhibition, with 250 nM of torin 1 (Selleck Chemicals S2827) or DMSO as the vehicle control for the same periods of time. Autophagic flux was assessed by exposing cells to BafA1 (100nM) for 2h.

### Florescence microscopy

For all cell culture microscopy experiments, HeLa cells and cortical neurons were cultured on Nunc Lab-Tek II 4 or 8-well glass chamber slides (Thermo Fisher Scientific). After exposure to autophagy inducing conditions, cells were fixed for 7 minutes with 4% PFA in PBS at room temperature and washed three times with PBS. For detection of GFP-LC3 or mCherry-capsid, a coverslip was mounted onto each slide with VECTASHIELD Antifade Mounting Medium with DAPI (Vector Laboratories H-1200), the perimeter of the coverslip was sealed with nail polish and allowed to dry overnight at room temperature before imaging. For immunofluorescence, both HeLa cells and cortical neurons were permeabilized after PFA fixation with 100% methanol chilled at −20℃ for 20 minutes followed by 1 h blocking with PBS containing 3% BSA (Sigma-Aldrich; A9418). Cells were stained with antibodies against SIN capsid, SQSTMI/p62 and phospho-SQSTM1/p62 for 2 h at room temperature in blocking buffer. After three washes, cells were treated with Alexa Fluor secondary antibodies for 1 h at room temperature, washed three times, and then mounted. All fluorescence imaging was performed using a Zeiss AxioImager Z2 microscope equipped with a Photometrics CoolSnap HQ2 CCD camera using a Zeiss PLAN APOCHROMAT 20X/0.8 NA wide-field objective or PLAN APOCHROMAT 40X/0.9 NA oil immersion objective. In all experiments a secondary antibody control was used for thresholding the background. Images were taken randomly of areas with similar cell density with the experimenter blinded to the experimental condition. HeLa cells SINV capsid/p62 and phospho-p62/total p62 images were taken in 0.3um Z-stacks and deconvoluted with AutoQuant X3 software. Image analysis of the p62 inclusion bodies was performed using Fiji software through a custom-written macro.

### SINV growth curves

To perform high MOI or low MOI multi-step viral growth curve analysis, HeLa cells were cultured in triplicate on 6-well plates and infected with either MOI of 10 or 0.01, respectively for 1.5 h. Virus infection media was replaced with fresh culture media. One hundred microliter samples were drawn from each well at specified time intervals and virus titers determined through Vero cell plaque assays.

### Cell death assay

For measurement of virus-induced cell death, cells were mock-infected or infected with SINV (MOI 10) for 24 h and then processed using the CellTiter-Glo Luminescent Cell Viability Assay, per manufacturer’s instructions (Promega, G7570). Luminescence was detected using a CLARIOstar Plate Reader (BMG Labtech).

### Co-immunoprecipitation

To investigate the interactions between CDKL5, p62 and SINV capsid, 10^7^ HeLa cells were cultured on 150 mm plates and either mock-infected or infected for 7h at MOI of 10 with recombinant SINV expressing HA-capsid. HeLa cells were scrapped from the plate and lysed in ice-cold lysis buffer (50 mM Tris-HCL (pH 7.5), 150 mM NaCl, 1 mM EDTA, 1% Triton X-100) containing complete proteinase inhibitor cocktail (Roche) and Halt phosphatase inhibitor cocktail (Thermo Fisher Scientific) for 30 min on ice. Cellular debris was removed with centrifugation for 10 minutes at 15,000 g at 4℃. Supernatants were pre-cleared with either dynabeads protein G (Thermo Fisher; 10003D) or protein G PLUS agarose beads (SCBT, sc-2002) for 1.5h at 4℃ with gentle agitation. After pelleting the beads, supernatants were mixed with either 0.8 μg/mL of rat anti HA antibody and 30 μL of pre-washed dynabeads protein G (Thermo Fisher; 10003D) or 40ul of pre-washed anti-Flag M2 affinity gel (Millipore Sigma; A2220) and then incubated at 4℃ overnight with gentle agitation. Dynabeads were pelleted using the DynaMag-2 Magnet (Thermo Fisher, 12321D) and anti-Flag beads by centrifugation at 3000g for 1 minute followed by 3 washes with ice-cold lysis buffer and two additional washes with lysis buffer containing 300mM NaCl. Immunoprecipitated proteins were eluted by adding 2X Laemmli sample buffer (Bio-Rad Laboratories) containing 5% β-mercaptoethanol (Bio-Rad Laboratories) and boiling sample for 7 minutes followed by western blot analysis.

### Protein lysate preparation and Western blot analyses

HeLa cells and cortical neurons were lysed with ice-cold 1X RIPA buffer (Cell Signaling Technology, 9806) supplemented with proteinase inhibitor cocktail (Sigma) and Halt phosphatase inhibitor cocktail (Thermo Fisher Scientific) and incubated on ice for 30 min. The insoluble components of the lysates were pelleted by centrifugation at 15,000 g at 4℃ for 15 min, and the residual supernatants mixed with 2X Laemmli sample buffer containing 5% β-mercaptoethanol. For analysis of p62 in total lysates, cells were directly lysed in 2x Laemmli sample buffer, boiled for 3 minutes in a 100℃ heating block, and then sonicated in an ultrasonic bath for one minute. For protein analysis in the soluble versus insoluble pellet, the pellet was washed with 1X RIPA buffer and resuspended with 2X Laemmli sample buffer at a ratio of 3:1. Proteins were separated on gradient 4-20% Mini-PROTEAN TGX precast protein gels (Bio-Rad Laboratories) and transferred onto PVDF membranes (Bio-Rad Laboratories). For blocking, membranes were incubated in 5% non-fat milk in either 1X TBS or PBS containing 0.01% Tween 20. Primary and secondary antibodies were diluted in blocking buffer. Proteins were detected with SuperSignal West Pico PLUS Chemiluminescent Substrate (Thermo Fisher Scientific) using a Bio-Rad Chemidoc imager. Band densitometry was determined through Fiji software.

### Flow Cytometry

WT and CDKL5 KO HeLa cells infected with SINV/mCherry-capsid were dissociated with 0.25% trypsin, fixed with 4% PFA for 5 minutes and resuspended in PBS with 3% BSA. Cells positive for mCherry-Capsid were detected through an LSRII-HTS flow cytometer (BD Biosciences) and data analyzed using FlowJo software (Version 9).

### Quantitative real-time PCR (qRT-PCR)

To assess p62 expression, RNA from HeLa cells infected with SINV was extracted at indicated time points using RNeasy Plus Mini Kit (Qiagen) and reverse-transcribed using the iScript cDNA Synthesis Kit (Bio-Rad Laboratories; 1708891). QuantiFast SYBR Green RT-PCR Kit (Qiagen, 204156) was used for the qPCR reaction and measurement performed on the 7500 Fast Real-Time PCR System (Applied Biosystems). The following primers were used for the reaction: p62 forward primer: 5’-TACGACTTGTGTAGCGTCTG-3’, p62 reverse primer: 5’-CGTGTTTCACCTTCCGGAG-3’, GAPDH forward primer: 5’-CGTGTCAGTGGTGGACCTG-3”, GAPDH reverse primer: 5’-CGTCAAAGGTGGAGGAGTGG-3.’

### In vitro Kinase Assays

For the reaction, 0.3 μg of recombinant human CDKL5 (1-498) (ThermoFisher, A33353) was combined with 3 μg of recombinant human SQSTM1/p62 protein (GeneTex, GTX68012-pro) in buffer containing 20 mM HEPES (pH 7.5) and 10 mM MgCl_2_ in the presence or absence of 50 μM [γ-P] ATP. The reaction was incubated at 30℃ for up to 45 minutes, stopped by adding 4x Laemmli sample buffer, then subjected to SDS-PAGE for Coomassie staining and radioactivity detection on x-ray film. This same reaction was also done using recombinant CDKL5 or TBK-1 (positive control) combined with recombinant p62 protein, with or without 100 μM non-radioactive ATP, and phosphorylation of p62 detected by western blot using phosphospecific antibodies. To compare the ability of WT and KD CDKL5 to phosphorylate recombinant p62 protein at Thr269/Ser272, HeLa cells stably expressing EV, WT or KD CDKL5 were lysed in ice-cold buffer (50 mM HEPES (pH 7.5), 300 mM NaCl, 1% Triton X-100) supplemented with proteinase inhibitor cocktail (Sigma) and Halt phosphatase inhibitor cocktail (Thermo Fisher Scientific). Pull down of Flag was performed as described above with anti-Flag M2 affinity gel. The beads were washed five times with lysis buffer containing 0.5 M NaCl and then three times with the in vitro kinase buffer (20 mM HEPES (pH 7.5), 10 mM MgCl_2,_ and 1 mM DTT) before resuspending the beads in 20 μl of in vitro kinase buffer with 100 μM ATP. The reaction was incubated at 30℃ for 45 minutes. Phosphorylation analysis was performed using Western blot.

### Viral infection of mice

Mice were infected intracerebrally into the right cerebral hemisphere with SINV and HSV-1 and subcutaneously with CHIKV. Viruses were diluted in Hank’s Balanced Salt Solution (HBSS, Thermo Fisher Scientific) and 30μL inoculum was used per mouse. For SINV, the dsTE12Q strain was used to infect seven-day-old *CDKL5*^WT^ and *CDKL5*^KO^ neonates with 1×10^3^ pfu. Separately, seven-day-old neonates were infected with 1×10^5^ pfu of CHIKV strain 06-02. For HSV-1 infection, 5×10^4^ pfu of HSV-1ΔBBD strain was used to infect anesthetized eight- to ten-week-old littermate *CDKL5*^WT^ and *CDKL5*^KO^ mice. For mortality studies, mice were monitored daily for 21 days. To perform brain tissue analysis in SINV infected mice, mice from each genotype were randomly selected for dissection at days 1, 4 and 7. The right hemisphere was snap frozen in liquid nitrogen, homogenized in HBSS and used for plaque assay titration to determine the viral load. The left hemisphere was fixed in 4% PFA, cryoprotected in 30% sucrose and embedded in paraffin for histology studies.

### Histology

For in vivo cell death analysis, terminal deoxynucleotidyl transferase dUTP nick end labeling (TUNEL) staining of paraffin-embedded sagittal sections was performed using Apoptag Peroxidase In Situ Apoptosis Detection Kit (EMD Millipore, S7100), per the manufacturers’ instructions. For immunohistochemistry, brain sections were stained with rabbit anti-SINV capsid antibody (1:5000 dilution) and the Vectastain Elite ABC HRP kit (Vector Laboratories, PK-6101) was used for signal detection, per manufacturer’s instructions. To capture the entire brain section, a Zeiss Axio Scan.Z1 slide scanner equipped with a Zeiss PLAN APOCHROMAT 20X/0.8 NA objective (Carl Zeiss Microscopy, UT Southwestern Whole Brain core facility) was used image each brain section. The number of TUNEL-positive or capsid positive cells per mouse brain sections was counted by an observer blinded to the condition and genotype.

### Statistical analyses

All statistical analyses were performed using GraphPad Prism Software (version 9). The individual statistical tests used are indicated on the figure legends. For in vitro studies, data was analyzed using unpaired two-tailed *t*-test when comparing two groups and Analysis of Variance (ANOVA) when comparing more than two groups, all under the assumption of normality. Mann-Whitney U test was used for a nonparametric comparison between two groups of mice when analyzing in vivo pathogenesis data. For mouse mortality studies, Kaplan-Meyer survival curves were generated and analyzed through log-rank tests to determine statistical significance.

## Acknowledgements

We dedicate this article to the memory and legacy of Dr. Beth Levine whose intellectual and financial contributions were fundamental to this work. This work was supported by NIH U19 AI142784 (B.L., M.U.S., T.A.R.), 5T32DK007745-23 (E.B.), K08AI163377 (J.W.T.), and The Welch Foundation I-1927-20200401 (J.L.J). We thank Diane Griffin for the capsid antibody, Deborah J. Lenschow for the CHIKV strain, Herbert W. Virgin and Robert Orchard for the ATG7 KO HeLa cells, Noboru Mizushima for the GFP-LC3 mice, and Lori Nguyen for assistance with animal experiments. The authors would like to acknowledge the assistance of the UT Southwestern Quantitative Light Microscopy Core, a Shared Resource of the Harold C. Simmons Cancer Center, supported in part by an NCI Cancer Center Support Grant, 1P30 CA142543-01. Many thanks to H.W. Virgin, Lynda Bennett, Xiaonan Dong and Yang Liu for helpful discussions on experiments. Tiffany Reese is the W. W. Caruth, Jr. Scholar in Biomedical Research and Michael Shiloh acknowledges the support of the Disease Oriented Clinical Scholars Program, both at University of Texas Southwestern. The authors would also like to thank Linda W. and Milledge A. Hart III for their generous support of autophagy research.

## Authors contributions

Conceptualization, J.W.T., B.L., J.L.J., J.K.P., T.A.R., M.U.S.; Formal analysis, J.W.T., T.A.R., M.U.S.; Funding acquisition, J.W.T., B.L., T.A.R.; Investigation, J.W.T., Z.Z., S.S., K.H., Y.W., V.S., G.U.; Project administration, J.W.T., M.U.S., T.A.R.; Supervision, J.W.T., B.L., J.L.J., J.K.P., T.A.R., M.U.S.; Visualization, J.W.T., T.A.R., M.U.S.; Writing – original draft, J.W.T., J.K.P., T.A.R., M.U.S.; Writing – review & editing, All authors.

## Supplemental Figure Legends

**Supplemental Figure 1.**
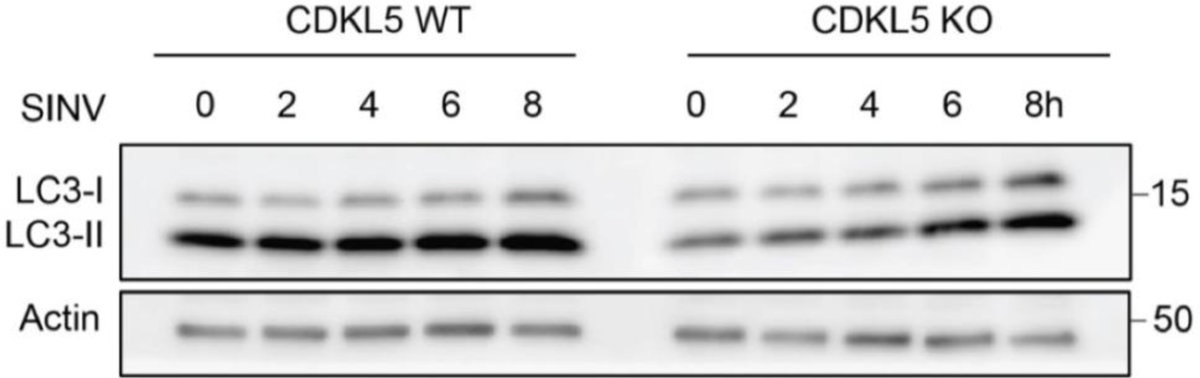
CDKL5-deficient cells have decreased autophagy at baseline and post SINV infection. LC3-I and LC3-II Western blot of WT and CDKL5 KO HeLa cells infected with SINV (MOI 10) over a time course. Blot representative of three independent experiments.

**Supplemental Figure 2.**
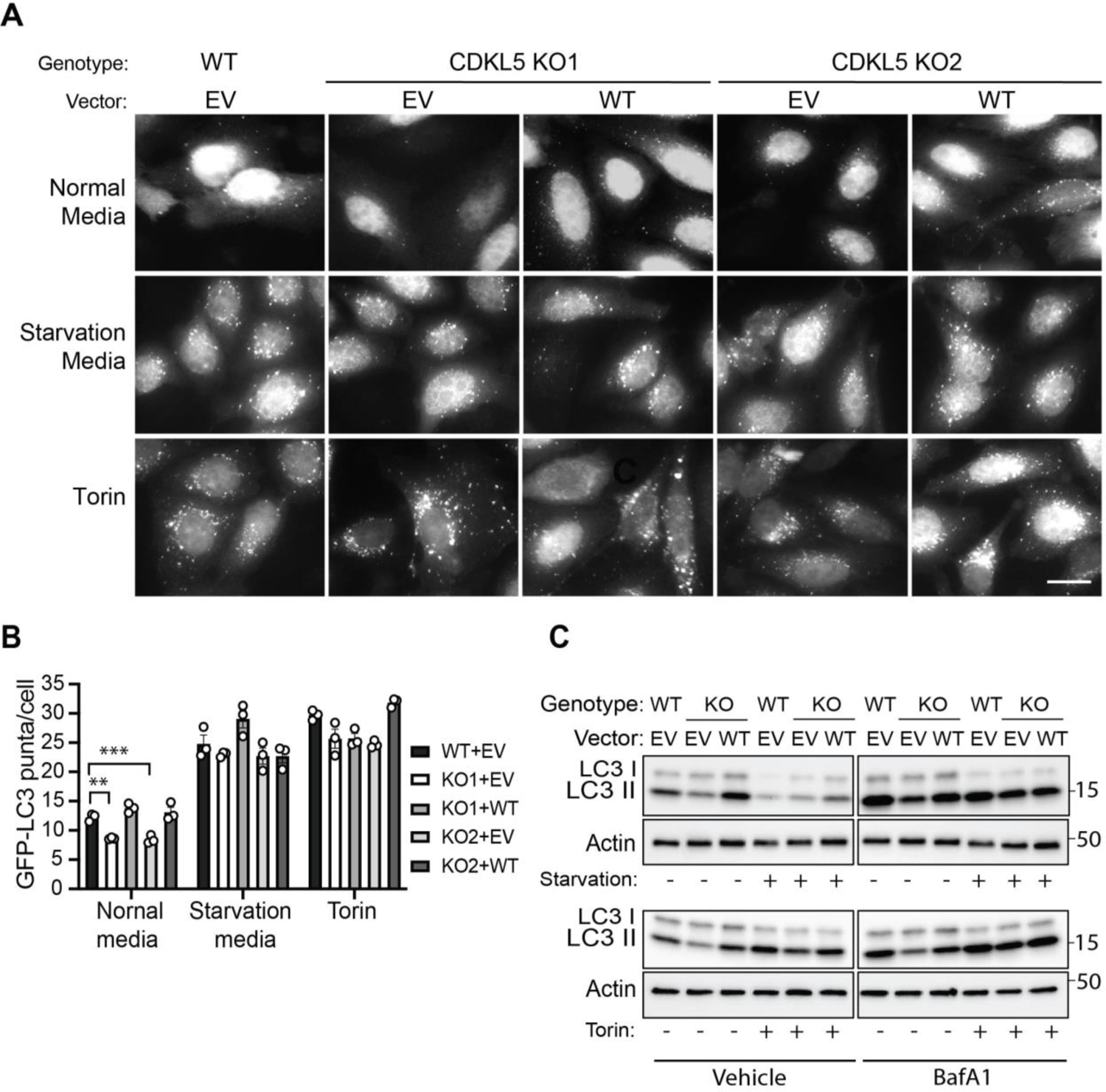
CDKL5 does not impact torin or amino-acid starvation induced autophagy. (A,B) WT and CDKL5 KO/GFP-LC3 HeLa clones reconstituted with EV or WT CDKL5 were treated with normal media, EBSS (starvation) medium or Torin (250 nM) for 1h. (A) Representative fluorescent micrographs of GFP-LC3 puncta. Scale bars = 20 μm. (B) Quantification of puncta per cell. Bars represent mean ± SEM of triplicate samples with at least 100 cells analyzed per sample. Similar results were seen in three independent experiments. Statistical analysis by one-way ANOVA with Dunnett’s test for multiple comparisons (\*\**p* < 0.01 and \*\*\**p* < 0.01). (C) Western blot of LC3-I and LC3-II from WT and CDKL5 KO HeLa cells reconstituted with WT CDKL5 or EV and treated with normal media, EBSS medium or Torin for 3h in the presence or absence of BafA1 (100uM) for the final 2 hours. Blot representative of 3 independent experiments.

**Supplemental Figure 3.**
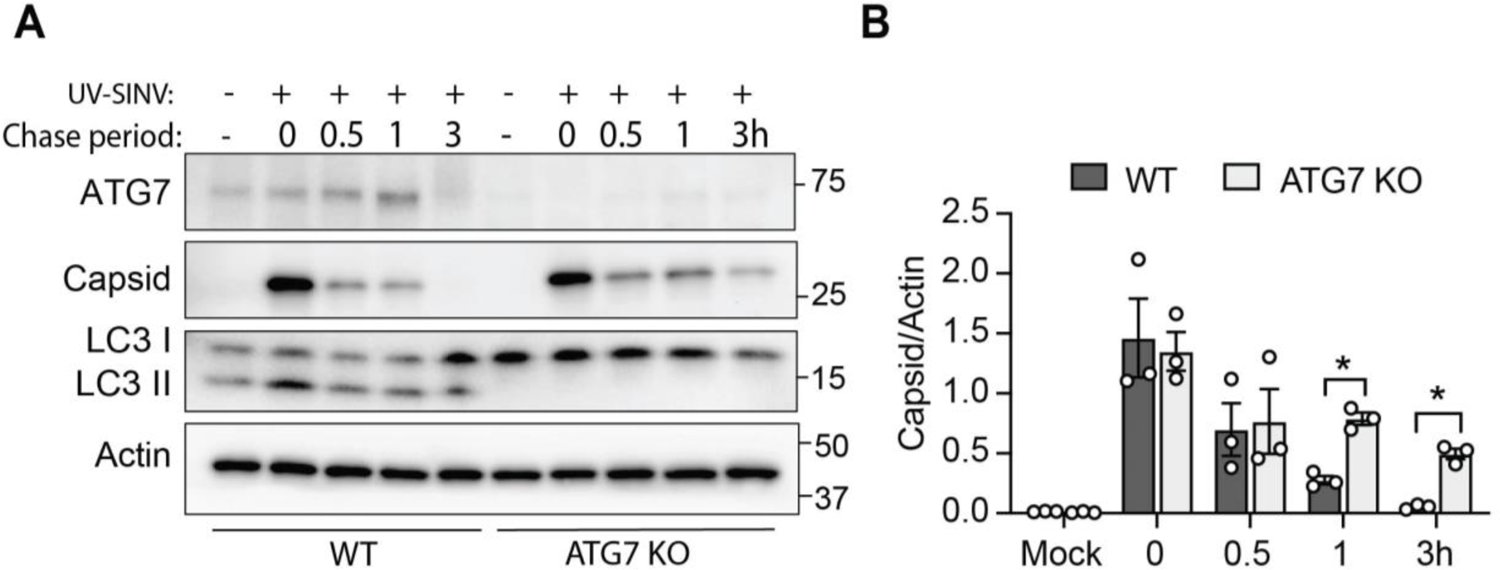
ATG7 KO cells are defective for capsid clearance after infection with UV-inactivated virus (A,B) WT and ATG 7 KO cells were exposed to UV-inactivated SINV (equivalent to 500 MOI) for 1 h, washed, and then lysates harvested at indicated time points. (A) Western blot of ATG7, capsid and LC3I/LC3II and (B) quantification of capsid/actin ratio by densitometry. Bars are mean ± SEM from three independent experiments. *p* values were determined by one-way ANOVA with Sidak’s multiple comparison test (\**p* < 0.05).

**Supplemental Figure 4.**
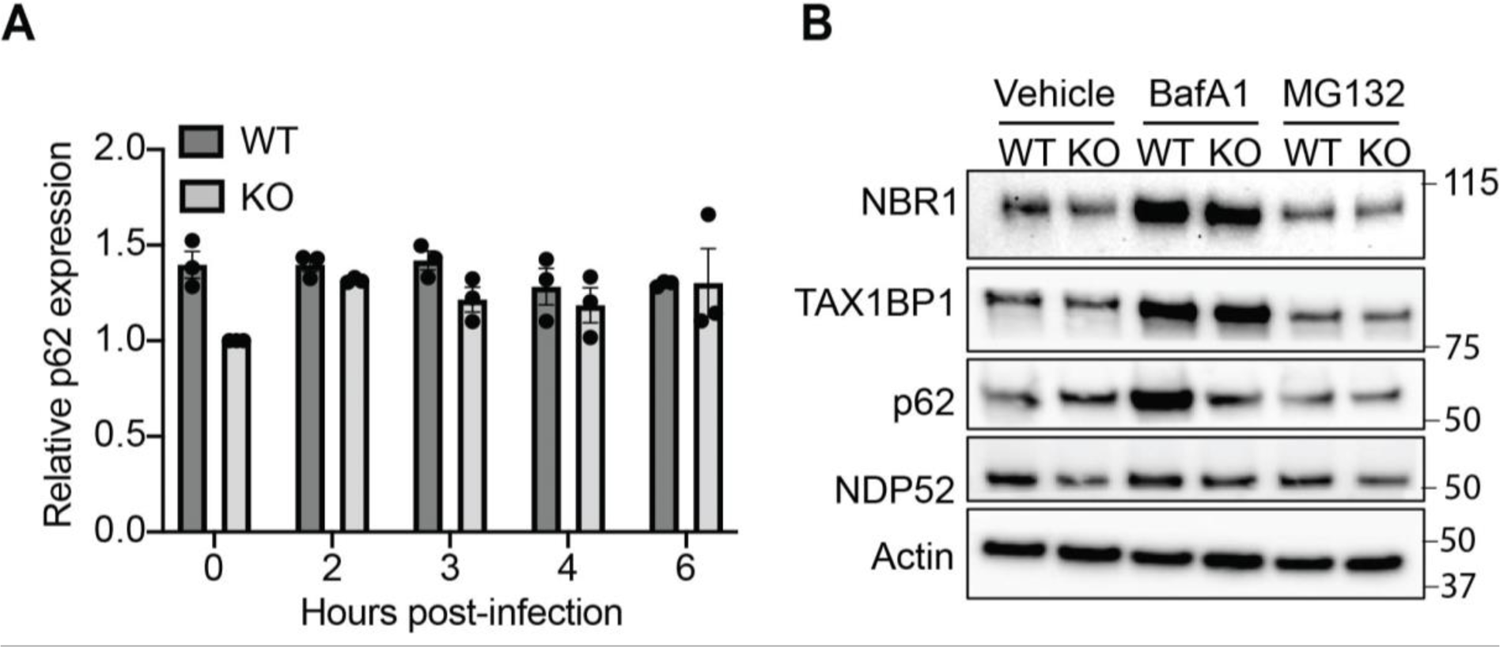
p62 expression and selective autophagy protein levels (A) *p62* expression in WT and CDKL5 KO HeLa cells infected with SINV (MOI = 10) was determined by RT-qPCR normalized to GAPDH. Bars mean ± SEM of triplicate samples and no significance noted on comparisons by unpaired two-tailed t-test; data representative of 3 independent experiments. (B) Western blot of autophagy receptors in WT and CDKL5 KO HeLa cells treated with DMSO (vehicle), BafA1 (100 μM, 4h) and the proteosome inhibitor MG132 (10 μM, 8h). Data are representative of two independent experiments.

**Supplemental Figure 5.**
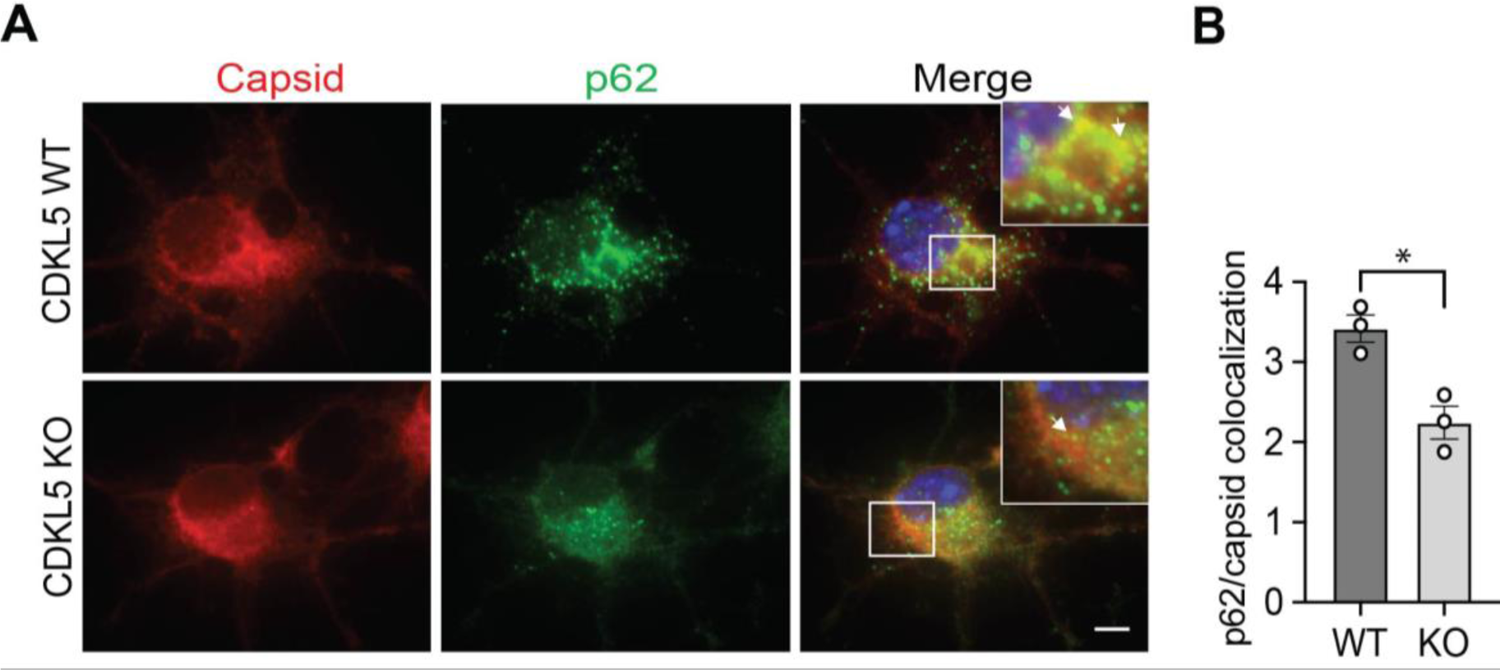
CDKL5-deficient cortical neurons have defective capsid and p62 interaction (A,B) Primary cortical neurons isolated from littermate *CDKL5*^WT^ and *CDKL5*^KO^ mice and cultured for 7 days were infected with SINV/mCherry-capsid virus (MOI = 10, 8h) then analyzed through immunofluorescence for mCherry-capsid (red) and p62 (green) colocalization. (A) Representative fluorescent micrographs. Arrows denote mCherry-capsid^+^/p62^+^ puncta. Scale bar 10 μm. (B) Quantification of mCherry-capsid and p62 positive puncta. Bars are mean ± SEM from neurons isolated from 3 independent embryos per genotype; at least 50 neurons per sample counted. *p* value was determined by unpaired two-tailed *t*-test (\**p* < 0.05).

**Supplemental Figure 6:**
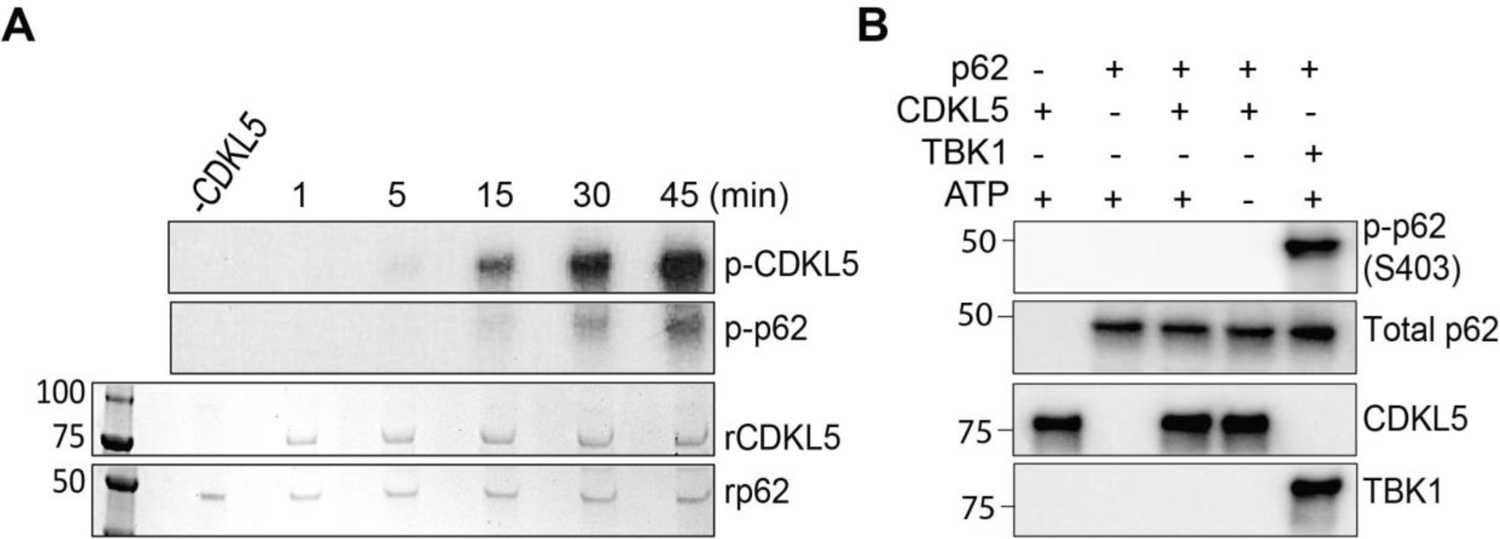
Kinetics of CDKL5 phosphorylaton of p62 by in vitro kinase assay (A) In vitro kinase assay with recombinant CDKL5 and p62 was performed using [^32^P] ATP and the reaction stopped at the indicated time points. The *Top* two panels are autophosphorylation of CDKL5 and phosphorylation of p62 detected by ^32^P-autoradiography. The *bottom* two panels show Coomassie Brilliant Blue staining of the gel. Blot representative of two independent experiments. (B) In vitro kinase assay with recombinant CDKL5 (1-498 aa) or TBK-1 and p62 using non-radioactive ATP and detection of p62 phosphorylation by western blot using anti-S403 phosphospecific antibody. Blot representative of two independent experiments.

